# Identification of small molecule inhibitors of *Trypanosoma* PEX15– PEX6 interaction

**DOI:** 10.1101/2025.10.03.680009

**Authors:** Lisa Hohnen, Bettina Tippler, Firat Tiris, Ralf Erdmann, Vishal C. Kalel

**Affiliations:** Department of Systems Biochemistry, Institute of Biochemistry and Pathobiochemistry, Faculty of Medicine, Ruhr-University Bochum, 44801 Bochum, Germany

**Keywords:** AlphaScreen, drug discovery, glycosomes, peroxin, *Trypanosoma*, PEX15, PEX6

## Abstract

Trypanosomatid parasites that cause life threatening tropical diseases harbor specialized essential organelles, called glycosomes. Like other peroxisome-related organelles, the biogenesis of glycosomes is mediated by proteins known as peroxins (PEX). A cascade of PEX protein-protein interactions (PPIs) is essential for glycosome function and parasite survival. Accordingly, small molecule inhibitors of PEX proteins that disrupt glycosomal matrix or membrane protein import have been reported as potential therapies for trypanosomiasis. We recently identified the long sought-after *Trypanosoma* PEX15 (*Tb*PEX15), which anchors the PEX1-PEX6 complex to the glycosomal membrane for recycling of the receptor PEX5. Defects in this process cause PEX5 degradation, mislocalization of glycosomal matrix proteins and parasite death. In this study, we targeted the interaction between *Tb*PEX6 and *Tb*PEX15. Recombinant *Tb*PEX6 and *Tb*PEX15 were purified, and their interaction was confirmed by *in vitro* pull-down assays and size exclusion chromatography. Furthermore, we established an AlphaScreen-based method to identify small molecule inhibitors of this PPI. Screening of a drug-repurposing library identified two inhibitors with trypanocidal activity against *T. brucei in vitro* and the amastigote stage of *T. cruzi*. Given its essentiality and low sequence similarity to its human homolog, parasite PEX15 and its interaction with PEX6 represent promising targets for the development of new therapies against trypanosomatid infections.

## Introduction

Insect transmitted trypanosomatid parasites cause three of the twenty-one neglected tropical diseases (NTDs). These diseases, specifically Human African Trypanosomiasis (HAT), also called African Sleeping Sickness, Chagas disease and Leishmaniasis are caused by *Trypanosoma brucei*, *T. cruzi* and several species of *Leishmania*, respectively. Current therapies, which are available to treat these diseases are limited in number, cause severe side effects and are ineffective due to emergence of drugs resistance. Thus, new drugs for effective treatment are of crucial importance. One way to target parasites for medical intervention is to disrupt biogenesis of their essential specialized peroxisome – the glycosome [1–5]. Glycosomes are multifunctional organelles, for which various matrix proteins are post-translationally imported. [6,7] Among others, glycosomes compartmentalize the first seven glycolytic enzymes [8,9]. This compartmentation is crucial for the parasite’s survival as the enzymes lack negative feedback regulation. Hence, upon mislocalization of these enzymes, turbo-glycolysis takes place, which leads to ATP depletion and finally cell death [10,11].

Glycosomal matrix protein import occurs through a transient import pore assembled by so-called peroxin (PEX) proteins. Proteins destined for the glycosomal matrix are recognized via their peroxisome targeting signal (PTS). Most cargo proteins carry a C-terminal PTS1 signal characterized by a tripeptide consensus sequence of [(SAC)(KRH)(LM)], with the preceding nine amino acids also influencing cargo-receptor binding [12]. These proteins are recognized by the cytosolic receptor PEX5. Less common are cargos with a PTS2 signal, which is close to the N-terminus and has a consensus sequence of R[L/V/I/Q]xx[L/V/I/H][L/S/G/A]x[H/Q][L/A] [13]. These are bound by the co-receptor PEX7, which then binds to PEX5 for glycosomal import. Upon binding of cargo-loaded PEX5 to the docking complex consisting of PEX14, PEX13.1 and PEX13. 2, a transient import pore forms, through which cargo protein is translocated into the glycosomal matrix. During or after cargo release, a complex of RING-finger peroxins ubiquitinate PEX5. This marks PEX5 for recycling upon monoubiquitination or for proteasomal degradation in case of polyubiquitination. Monoubiquitinated PEX5 is exported back to the cytosol by the heterohexameric AAA+ ATPase PEX1 and PEX6, which is brought to the membrane by Pex15p (yeast)/PEX26 (human)/APEM9 (plant).

It had already been shown that defects in Pex15p/PEX26/APEM9 result in disruption of peroxisomal protein import and pexophagy in other organisms [14,15]. In *H. sapiens* mutations in *PEX26* gene leads to different forms of peroxisome biogenesis disorders such as Zellweger syndrome, milder neonatal adrenoleukodystrophy (NALD) and infantile Refsum disease (IRD) [16–18]. Mutations in *PEX26* gene were also found to be associated with Heimler syndrome [19]. The cause for these disorders has been hypothesized to be polyubiquitinated PEX5-induced pexophagy due to a lack of functional AAA+ ATPase complex [20]. As peroxisomes have pro-tumorigenic functions [21], pexophagy induced by a reduction of PEX26 levels sensitizes drug-resistant cancer cells for therapy [22]. On the other hand, in colorectal cancer (CRC) *PEX26* is down-regulated and overexpression of PEX26 inhibits cell migration, invasion, and epithelial–mesenchymal transition (EMT) [23].

The suspected PEX15/PEX26 protein at the glycosomal membrane, had been long sought after. *Trypanosoma* PEX15 was recently identified in our proteomic analysis of purified glycosomal membranes [24]. Knock-down of protein expression of *Tb*PEX15 by RNAi leads to mislocalization of glycosomal matrix proteins and causes severe growth defects demonstrating its essentiality for parasite survival. Moreover, reduction of *Tb*PEX5 steady state levels upon *Tb*PEX15 RNAi suggests its involvement in the export of *Tb*PEX5, as recently also shown for *Tb*PEX1. Yeast two-hybrid (Y2H) assays confirm its interaction with *Tb*PEX6 and thus, its functional role in anchoring the *Tb*PEX1-*Tb*PEX6 complex to the glycosomal membrane. Since the interaction between PEX15 and PEX6 is crucial and *Tb*PEX15 shares very low sequence similarity with its human counterpart PEX26, the *Tb*PEX6-*Tb*PEX15 protein-protein interaction is an attractive drug target. In this study, we demonstrate a direct interaction between *Tb*PEX6 and *Tb*PEX15 by *in vitro* pull-down assays with recombinant proteins. Furthermore, we established an AlphaScreen-based assay and identified inhibitors of the *Tb*PEX6-PEX15 interaction from a drug repurposing library. These inhibitors show trypanocidal activity against *T. brucei* and *T. cruzi* parasites, and no significant cytotoxicity for human cells. The established assay is a valuable tool to further screen larger compound libraries and extending the study to *Leishmania* counterparts, with final goal of developing new therapies for trypanosomiasis and leishmaniasis.

## Results

### Demonstration of a direct *Tb*PEX15-*Tb*PEX6 interaction *in vitro*

PEX15 is a glycosomal membrane anchor for the AAA^+^-ATPases PEX1–PEX6. Functional recruitment of the ATPases by PEX15 to the peroxisomal membrane has been shown to be crucial for glycosome biogenesis [24]. Blocking this interaction will disrupt functional matrix protein import and hence, cause lethal effects on trypanosomes. We recently showed that *Tb*PEX15 lacking the transmembrane region (321-360aa) interacts with full-length *Tb*PEX6 in Y2H assay [24]. To confirm these results and to demonstrate a direct interaction, recombinant histidine-tagged *Tb*PEX15 protein lacking the C-terminal transmembrane domain (His_6_-*Tb*PEX15ΔTM, residues 1-320) and GST-tagged full-length *Tb*PEX6 (GST-*Tb*PEX6) were affinity purified (**Fig. S1**) and tested for interaction in an *in vitro* pull-down assay (**Fig. 1**). *Tb*PEX15ΔTM was clearly retained on GSH beads in the presence of GST-tagged *Tb*PEX6 and was eluted upon incubation with reduced glutathione demonstrating a direct interaction between these two proteins (**Fig. 1A**). GST protein alone (control) was not able to bind *Tb*PEX15ΔTM (**Fig. 1B**).

**Figure 1.**
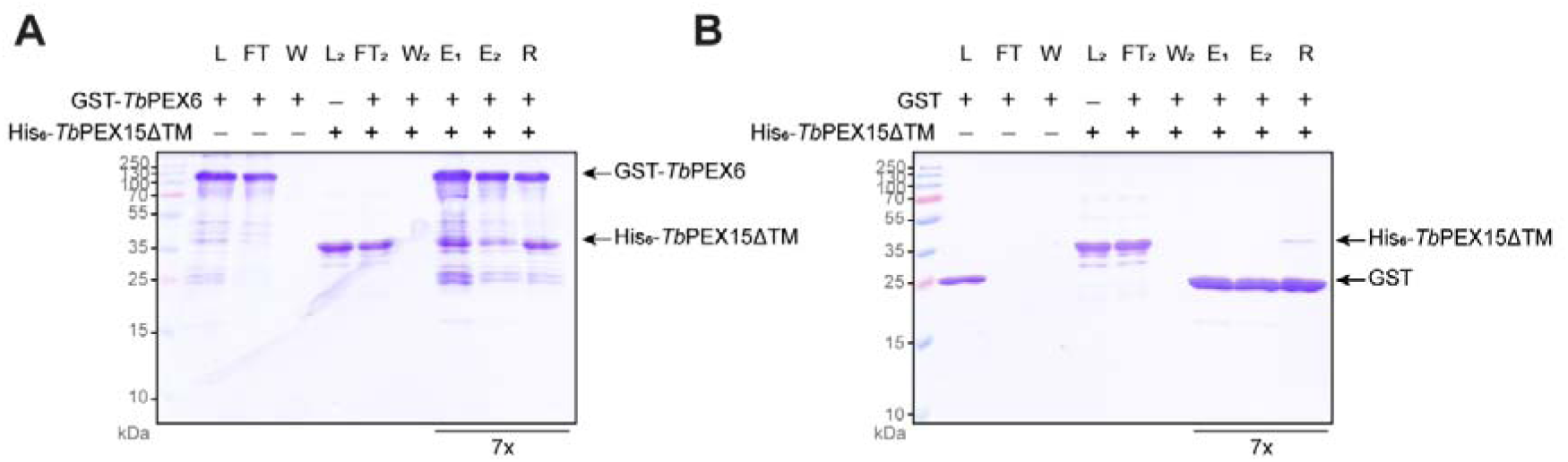
**Interaction between recombinant His_6_-*Tb*PEX15**Δ**TM (1-320aa) and GST-*Tb*PEX6 (full-length**). *In vitro* pull-down assay including GST as control protein. GST-*Tb*PEX6 (**A**) and GST (**B**) were bound to GSH agarose beads for 1 h (L). After removal of unbound protein by washing (W), the protein-bound beads were incubated with His_6_-*TbP*EX15ΔTM (1-320aa) protein for 1 hour (L_2_). Unbound protein is washed off (W_2_), followed by elution with reduced glutathione in wash buffer. After eluting twice (E_1_ and E_2_), the beads are heat denatured in 1x SDS-Laemmli buffer (R) to recover all protein still bound to the beads.

To investigate the stoichiometry of binding by size exclusion chromatography (SEC), we attempted removal of the GST-tag from *Tb*PEX6 as GST is known to dimerize and thus disturbs the analysis or produces artefacts. Removal of the GST-tag, however, destabilized the protein and resulted in aggregation of PEX6 (data not shown). Moreover, *Tb*PEX6 that had been tagged with a smaller His_6_ tag could not be overexpressed in a soluble state. We reasoned that *Tb*PEX6 may require a binding partner to retain its stability. Accordingly, we first pre-formed the GST-*Tb*PEX6 – *Tb*PEX15 complex on GSH beads, followed by cleavage of the GST-tag using PreScission protease. Indeed, we were able to remove the tag and retrieve a *Tb*PEX6 – *Tb*PEX15 complex. Interestingly, most of the tag-free *Tb*PEX6 was retained on the GSH beads and eluted only upon glutathione elution (**Fig. S2,** left panel; see **E_1_**). The soluble complex of tag-free *Tb*PEX6 and His_6_-*Tb*PEX15ΔTM was further separated by SEC, where both proteins co-eluted with a retention volume of 1.64 mL, corresponding to an apparent molecular mass of 161 kDa based on the calibration curve (**Fig. 2A**). Given that the theoretical molecular weight of the complex is 143.1 kDa, we conclude that *Tb*PEX6 and His_6_-*Tb*PEX15ΔTM interact in a monomeric state with 1:1 stoichiometry. *Tb*PEX15 also smeared and peaked around 2ml retention volume, which may represent its partially dissociated portion from *Tb*PEX6 during chromatography. GST-PreScission protease was also detected in the eluate, co-eluting in similar fractions of PEX6-PEX15 complex (**Fig. 2A**, bottom). However, this was attributed to GST dimerization, as the elution profile of GST-PreScission protease remained unchanged in the absence of tag-free *Tb*PEX6 or His_6_-*Tb*PEX15ΔTM (**Fig. 2B**). In addition, His_6_-*Tb*PEX15ΔTM alone also smears elutes in much later fractions (peak ∼1.8 - 2.0 mL), representing monomeric *Tb*PEX15.

**Figure 2.**
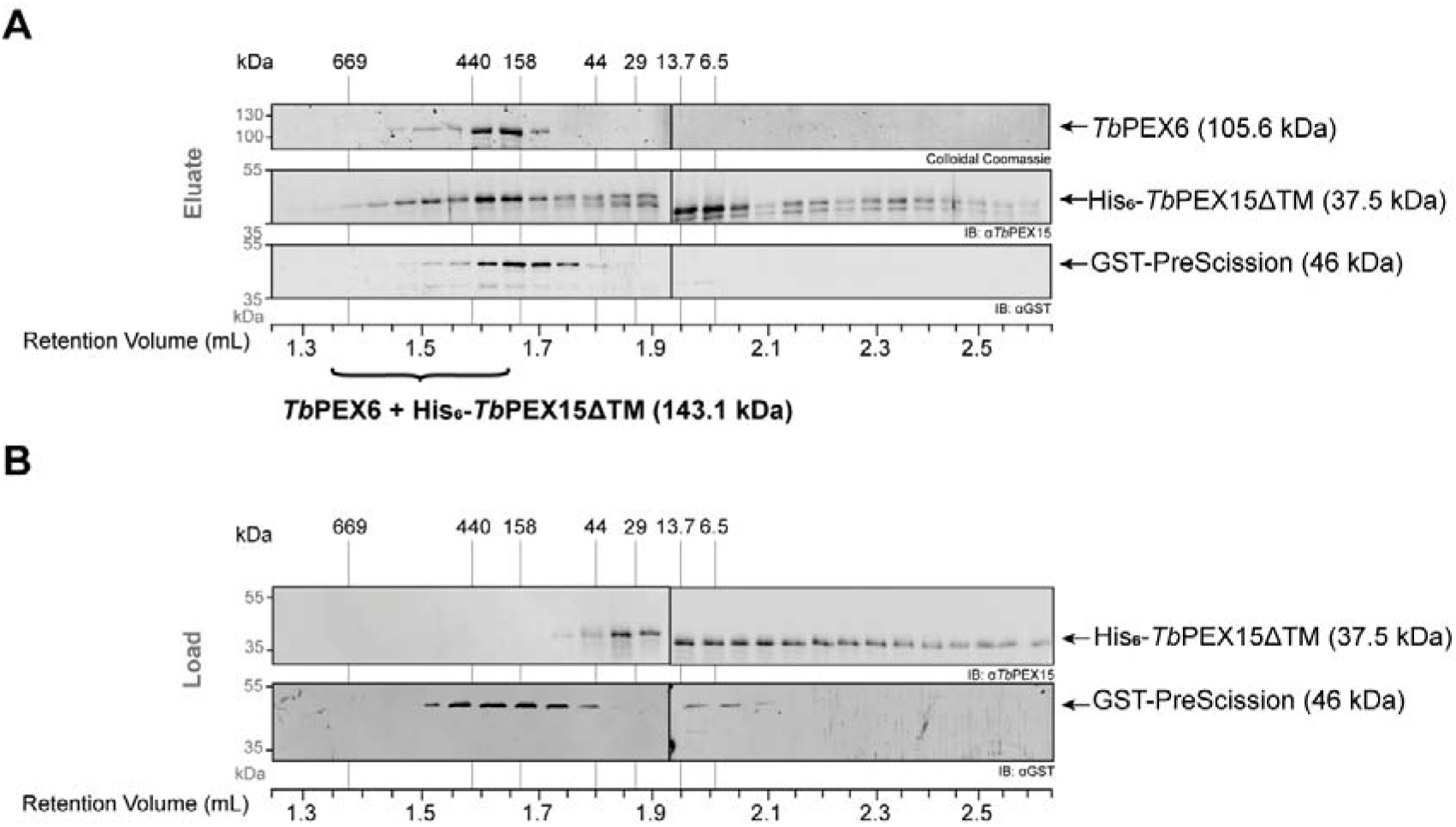
Analysis of *Tb*PEX15-*Tb*PEX6 complex by size-exclusion chromatography. **(A)** Size exclusion chromatography of the *Tb*PEX15-PEX6 complex was performed on a Superose™6 PC 3.2/30 column. Fractions were analyzed by SDS-PAGE followed by Colloidal Coomassie staining for *Tb*PEX6 (Top); and by or immunoblotting using α*Tb*PEX15 (middle) or αGST antibodies (bottom). His_6_-*Tb*PEX15ΔTM and *Tb*PEX6 eluted together with a retention volume of 1.64 mL corresponding to a molecular mass of 161 kDa. His_6_-*Tb*PEX15ΔTM was also detected in later fractions. GST-PreScission protease peaked with a retention time of 1.69 mL. This correlates to a molecular weight of 109.4 kDa, demonstrating dimerization of the fusion protein via the GST-tag. **(B)** The same retention time was monitored when GST-PreScission Protease alone was loaded onto the Superose™6 PC 3.2/30 column (lower panel). His_6_-*Tb*PEX15ΔTM alone is present in later fractions, peaking at a retention volume in range of 1.8-2.0 mL. The column was calibrated with thyroglobulin (669 kDa), ferritin (440 kDa), aldolase (158 kDa), ovalbumin (44 kDa), carbonic anhydrase (29 kDa), ribonuclease A (13.7 kDa) and aprotinin (6.5 kDa).

### Establishment of an AlphaScreen based *Tb*PEX15–PEX6 binding assay

To quantitatively analyze the *Tb*PEX15-PEX6 interaction, we established an *in vitro* Alpha (Amplified Luminescent Proximity Homogeneous Assay, PerkinElmer [25]) beads-based binding assay (**Fig. 3A**). The assay was established using same recombinant proteins i.e. His_6_-*Tb*PEX15ΔTM and GST-*Tb*PEX6, which were used for the *in vitro* pull down (**Fig. 1**). The optimal protein concentration for a robust Alpha signal was determined by cross-titration of both proteins with concentrations ranging from 0 to 300 nM (**Fig. S3**). Based on these results, we performed *vitro* binding assays in low nanomolar concentration range.

**Figure 3.**
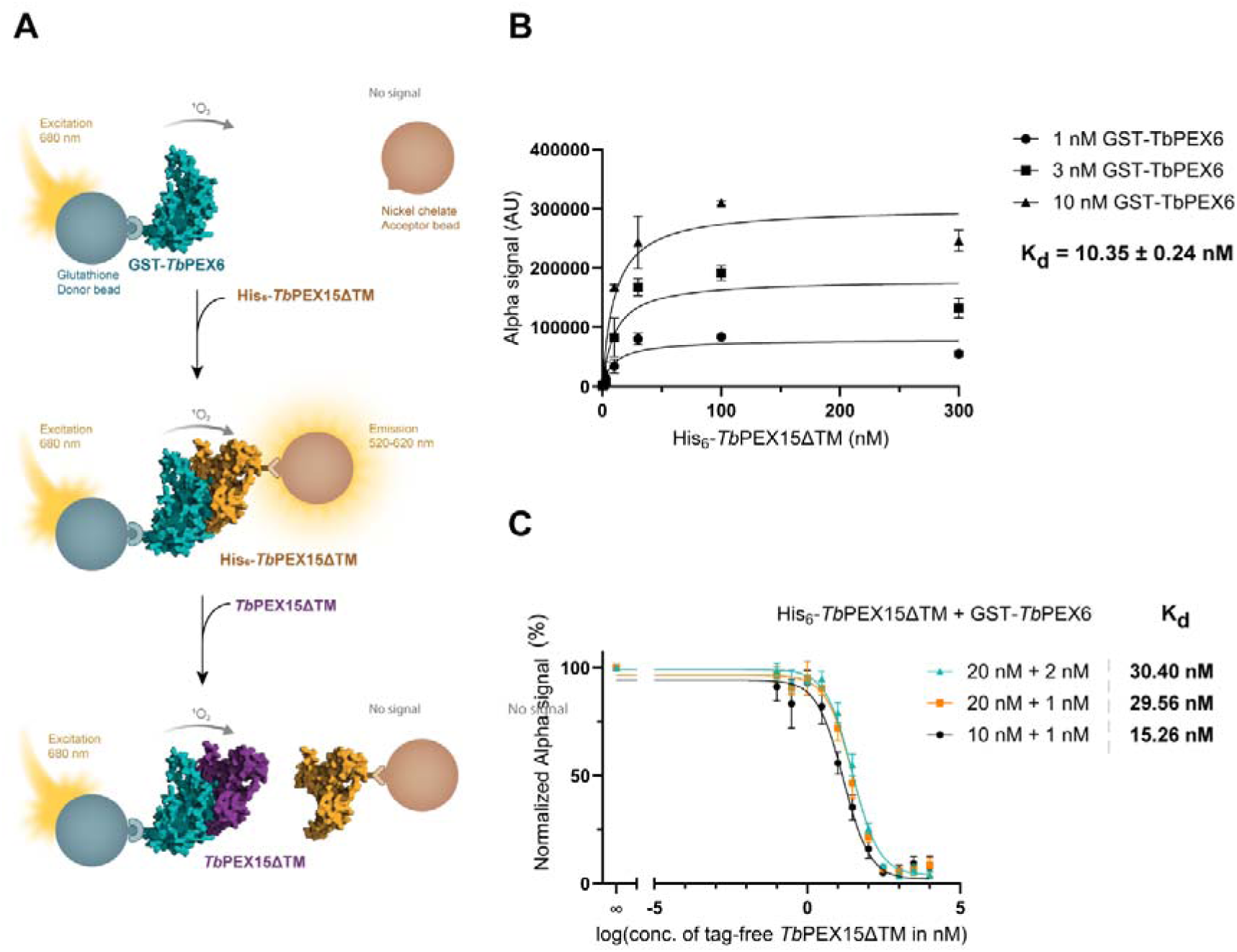
AlphaScreen-based *Tb*PEX15-PEX6 binding assay. Determination of apparent K_d_ by saturation and displacements assays using the AlphaScreen technology. **(A)** Schematic representation of the assay. GSH-donor and Ni-Chelate acceptor Alpha beads were used to perform assay. **(B)** His_6_-*Tb*PEX15ΔTM at different concentrations ranging from 30 nM to 300 μM was added to 1, 3 and 10 nM GST-*Tb*PEX6 as indicated. The results were collected from three independent experiments and analyzed using one site-specific binding model in GraphPad Prism 10.0.0. The dissociation constant (K_d_) was estimated to be 10.35 ± 0.24 nM for the three conditions. **(C)** Tag-free *Tb*PEX15ΔTM competes with His_6_-*Tb*PEX15ΔTM for interaction with GST-*Tb*PEX6. Increasing concentrations of Tag-free *Tb*PEX15ΔTM were titrated to a pre-incubated mixture of His_6_-*Tb*PEX15ΔTM and GST-*Tb*PEX6, resulting in a progressive decrease of the Alpha signal. Data points were collected from three replicates for each condition and analyzed using one site-fit K_i_ in GraphPad Prism 10.0.0. Signal was normalized to the positive control, which comprised both His_6_-*Tb*PEX15ΔTM and GST-*Tb*PEX6 without addition of tag-free *Tb*PEX15ΔTM. The K_d_ was calculated to be between 15.26 and 30.4 nM across the three conditions. Error bars represent the standard deviations.

First, we performed a saturation assay, where fixed concentrations of GST-*Tb*PEX6 was titrated with increasing concentrations of His_6_-*Tb*PEX15ΔTM (**Fig. 1B**). This revealed an apparent dissociation constant (K_d_ value) of 10.35 ± 0.24 nM. Next, we utilized displacement assays, where untagged *Tb*PEX15ΔTM was added to defined concentrations of GST-*Tb*PEX6 and His_6_-*Tb*PEX15ΔTM (**Fig. 1C**). The untagged protein competitively displaces the tagged binding partner, resulting in reduction in Alpha signal. The K_d_ determined from this assay ranged from 15.26 to 30.40 nM depending on the concentration of His_6_-*Tb*PEX15ΔTM, confirming a strong interaction between recombinant *Tb*PEX15 and *Tb*PEX6. According to the manufacturer’s guidelines, using lower protein concentrations in the assay provide a more accurate approximation of the apparent K_d_ in the displacement experiments. Thus, the value of 15.26 nM may represent the more reliable estimate.

### High-Throughput Screening of *Tb*PEX15–PEX6 Interaction Inhibitors using AlphaScreen™

AlphaScreen assays enable proximity-based PPI detection without the need for wash steps. Moreover, the assay setup can be miniaturized for 96, 384 or 1584 well plates for high-throughput screening of molecules that inhibit the specific PPI (**Fig. 4A**). This technology has already been used in several drug discovery campaigns for inhibitors targeting PEX protein interactions [1,2]. In this study, we established and optimized the AlphaScreen assay for inhibitor screening of the *Tb*PEX15-PEX6 interaction in a 384-well plate format with 25 µl reaction volume. According to the cross-titration results (**Fig. S3**) that provide robust and optimal Alpha signal, we used both proteins in a concentration of 10 nM. Each screening plate included various in-plate controls. (i) Wells without test protein served as negative control indicating the magnitude of background noise. (ii) The positive control comprised all components of the assay except any chemical compound showing maximum signal without compound interference. (iii) Addition of 1 M NaCl results in the dissociation of the *Tb*PEX15– PEX6 complex and was applied as an additional negative control [26]. In this study, we screened compounds from a drug repurposing library (MCE-HY-L035P-30 PartA) that contains FDA-approved drugs and compounds, which are currently in clinical trials for inhibition of *Tb*PEX15-PEX6 PPI (primary screen). Per plate, 320 of these compounds were screened, with the remaining wells containing the above-mentioned controls.

**Figure 4.**
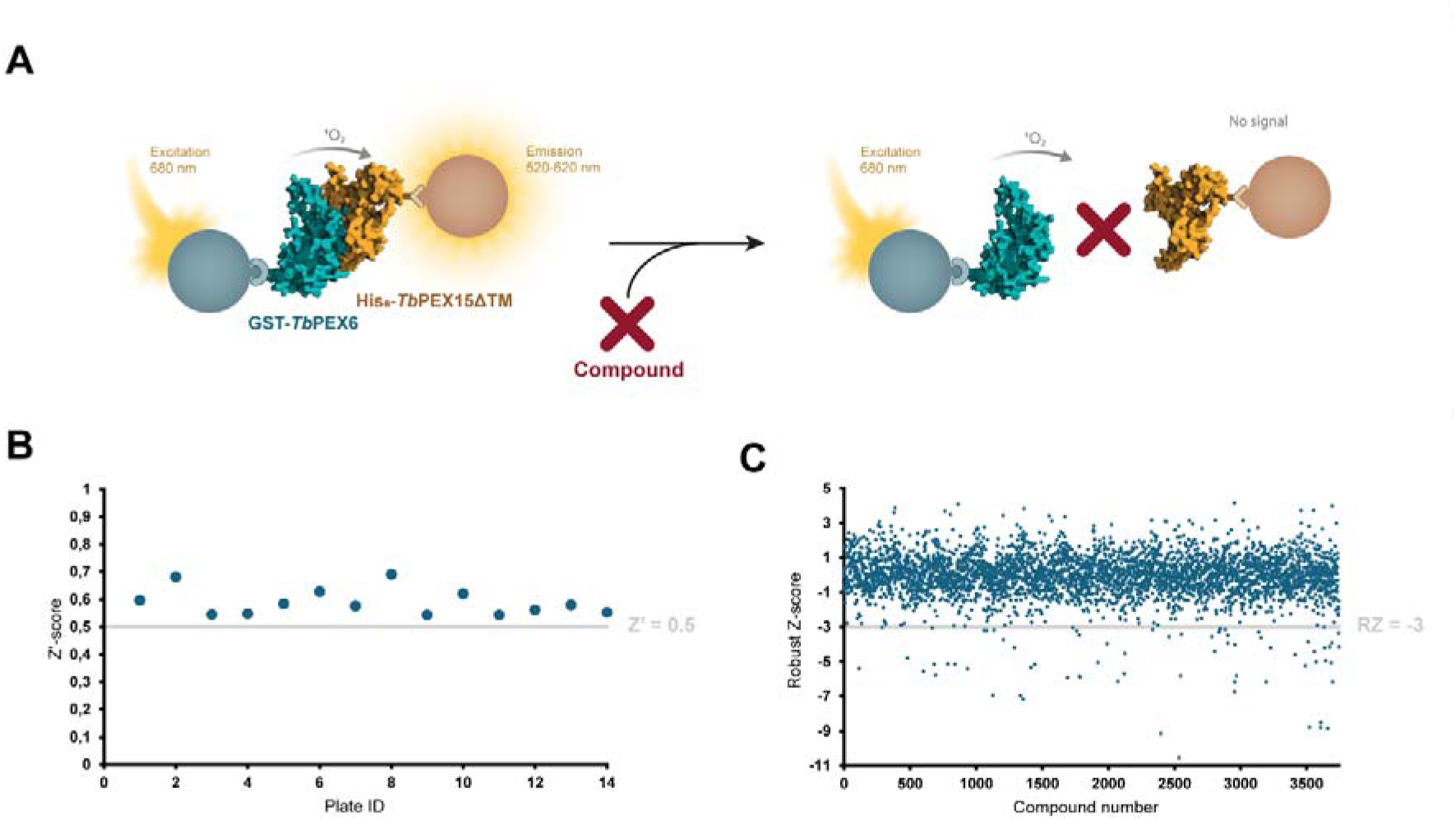
Establishment of the AlphaScreen-based High-Throughput assay for identification of *Tb*PEX6-PEX15 interaction inhibitors. (**A**) Alpha-technology is a bead-based system. The tagged proteins of interest bind to differently modified beads: donor and acceptor beads. The donor bead is excited at 680 nm and emits a singlet oxygen. Due to the short lifetime of singlet oxygen, energy transfer to the acceptor bead occurs only if the donor and acceptor beads are within a range of 200 nm to each other. A light signal is emitted by acceptor beads which can be quantitatively detected as Alpha signal between 520-620 nm. In the presence of molecules that inhibit the interaction between the proteins, the beads do not get into close enough proximity to each other and hence, the signal is abolished or reduced. (**B**) Z’-scores calculated for the 14 plates that were screened in this study. The Z’-score is >0.5 indicating the assay is robust and the result reliable. Within the 14 plates total of 3744 drugs were screened. (**C**) The robust Z-score was calculated for hit selection. Compounds with a robust Z-score <-3 were selected as primary hits for counter screening.

To assess the reproducibility and validity of the assay, the Z’ factor was calculated for each screened plate [27]. A Z’ factor above 0.5 indicates that the results obtained for a specific plate are statistically reliable and can be considered for further analysis. In this study, 14 plates were screened for the primary screen with Z’ factors between 0.5 and 0.7 (**Fig. 4B**). These screens covered 3744 compounds that were tested for their ability to inhibit the *Tb*PEX15-PEX6 PPI at a single concentration of 50 μM/ 10 μM/ 15 μg/mL depending on the stock concentration of the parent drug. Next, we applied the robust Z score (RZ score) as an additional statistic criterion to shortlist hits from the primary screen. The RZ score, calculated for each tested compound using the median and median absolute deviation, is insensitive to outliers [28–30]. Initial hits were defined as compounds with an RZ score ≤-3 (**Fig. 4C**). Scores below this threshold indicate signals that are significantly reduced compared to the majority. Using this criterion, 71 compounds were selected.

The shortlisted 71 primary hits were further tested in a counter screen assay to detect false positive hits **(Fig. 5)**. False positive hits may interfere with the Alpha signal due to intrinsic fluorescence or quenching effects. Also, the compounds might bind unspecifically to either the affinity tags or the beads, thereby disrupting protein binding to the beads rather than the protein-protein interaction **(Fig. 5A**, schematic representation). Each compound or its respective solvent (H_2_O or DMSO) was incubated with GST-*Tb*PEX6 and His_6_-*Tb*PEX15ΔTM or GST-His_6_ in parallel. The signals were normalized to the positive control (no compound treatment) and displayed as relative inhibition levels. If the signal was decreased by ≥ 50% when incubated with tagged *Tb*PEX15–PEX6 and ≤ 30% if incubated with the GST-His_6_ fusion protein and the difference between both protein pairs was at least 50%, the compound was considered as a true and validated hit **(Fig. 5B)**. Based on this counter screen, six compounds were shortlisted **(Fig. 5B**, orange spheres) and procured as 10 mM stocks from MedChemExpress for further analysis.

**Figure 5.**
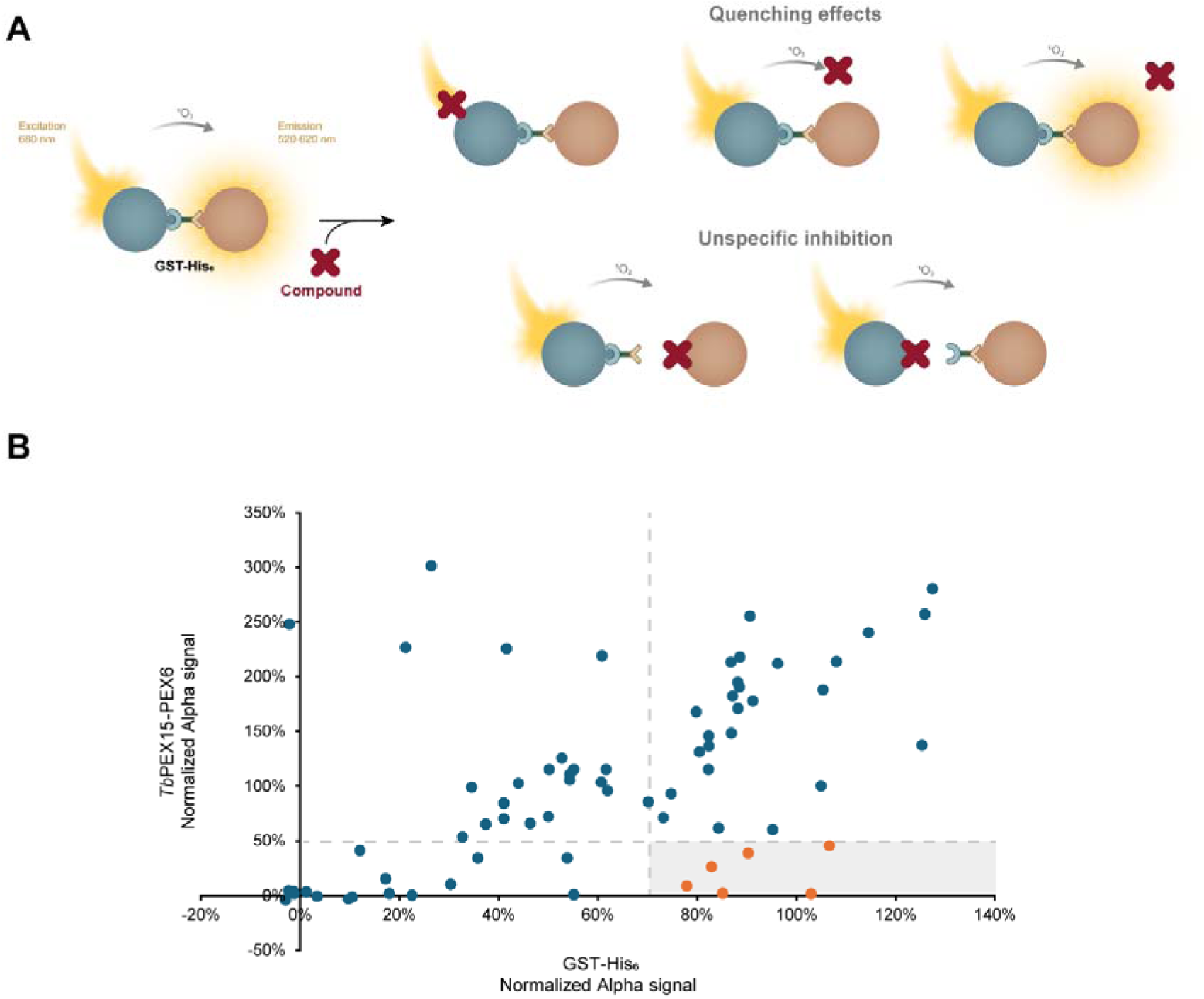
Counter screen for shortlisting true *Tb*PEX15-*Tb*PEX6 inhibitors. **(A)** Scheme depicting potential reasons for appearance of false positives and rationale of using GST-His_6_ as a counter screen protein. **(B)** Selected 71 primary hits were again screened against *Tb*PEX15ΔTM—*Tb*PEX6 (test screen) as well as GST-His*_6_* control protein (counter screen) to sort out false positives. Alpha signal normalized with the positive control (without compound treatment) for each primary hit from both screens were plotted. If a compound showed at least 50% inhibition (50% signal left) of *Tb*PEX15ΔTM—*Tb*PEX6 interaction and maximally 30% inhibition (70% signal left) in presence of GST-His*_6_* control protein, the compounds were classified as true hits and further evaluated.

The shortlisted true hit compounds are Compound X (CmpX, pseudo-named), the bisphosphonates clodronate and risedronate as well as the amino acid analog L-selenomethionine, the somatostatin analog vapreotide and the β-lactam-antibiotic ceftazidime. All the listed compounds are FDA-approved and sold to treat various diseases. We evaluated these compounds for their half maximal inhibitory concentrations (IC_50_) using dose-response AlphaScreen assays **(Fig. 6)**. Here, two compounds, CmpX and clodronate, had IC_50_ values in the nanomolar range – 102 nM and 199 nM (**Fig. 6** left panel, bottom and top). No IC_50_ value could be determined for risedronate (**Fig. 6** left panel, middle) as it appeared to interfere with the signal at higher compound concentrations. This interference was detected only in presence of the interacting protein pair, but not in their absence (data not shown). The IC_50_ of L-selenomethionine and vapreotide were determined as 3.06 μM and 18.73 μM, respectively (**Fig. 6** right panel, middle and top). For ceftazidime, no IC_50_ value could be calculated but it is expected to be >100 μM (Fig. 6 right panel, bottom).

**Figure 6.**
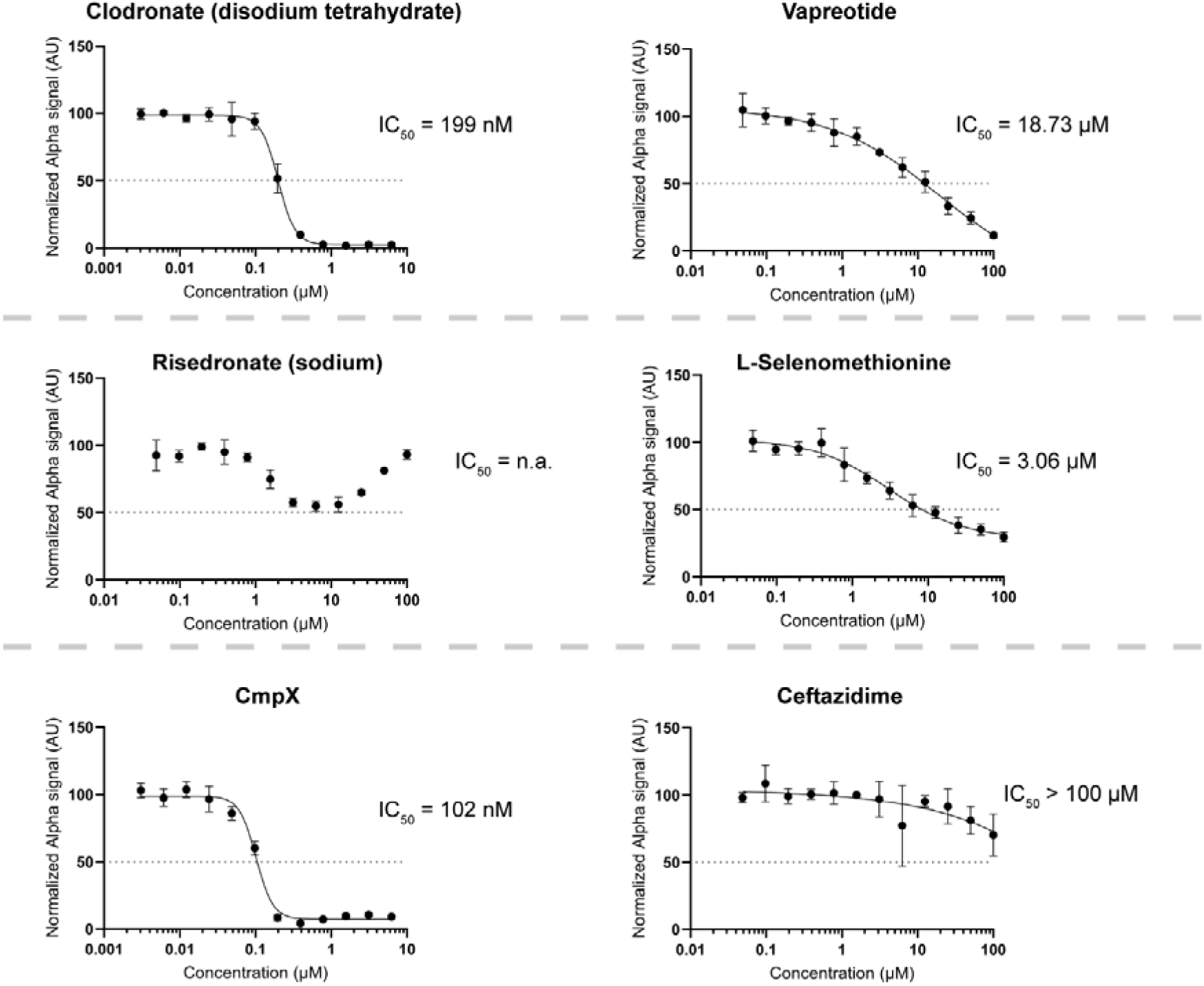
Dose-response curves of selected inhibitors in AlphaScreen based PPI inhibition assay. 6 compounds (clodronate, risedronate, CmpX, vapreotide, L-selenomethionine and ceftazidime) were tested in dose dependent manner for *Tb*PEX15-PEX6 interaction inhibition in Alpha assay. IC_50_ (concentration leading to 50% interaction inhibition) values of clodronate and CmpX are in the nanomolar range (199 nM and 102 nM). For risedronate, no IC_50_ value could be determined due to signal interference. Vapreotide and L-selenomethionine showed IC_50_ values of 18.73 μM and 3.06 μM, respectively. Ceftazidime shows an effect only at higher micromolar concentrations and the IC_50_ value is estimated to be above 100 μM. Shown are the results of three replicates, error bars indicate ±SD. IC_50_ values were determined via GraphPad Prism 10.0.0.

The identified compounds are reliable hits, since none of these compounds appeared in screens with *Leishmania* PEX3-PEX19 [4].

### Hit Validation

After the initial screen, the shortlisted compounds were also tested in an enzyme-linked immunosorbent assay (ELISA) to confirm their inhibitory effect in an orthogonal assay (**Fig. S4**). To this end, wells of a 96-well plate were coated with His_6_-*Tb*PEX15ΔTM. After blocking, compounds were added to the wells for a 30 min incubation at room temperature followed by addition of GST-*Tb*PEX6. After a total incubation time of 90 min, the solution was removed, and detection was performed utilizing TMB substrate and sulfuric acid. A high concentration of salt (1 M sodium chloride) solution was used as a positive control for disruption of the interaction. Cmp1, a compound which was identified from the same library to inhibit the *Ld*PEX3-*Ld*PEX19 interaction [4] but was not shortlisted in this study. Clodronate, risedronate and L-selenomethionine were confirmed to inhibit the interaction in a dose-dependent manner in the ELISA assay. CmpX and vapreotide interfered with signal detection, hence, could not be validated in this assay. Notably, ceftazidime, which showed only a weak inhibition in the AlphaScreen dose-response assay (**Fig. 6** left panel, bottom) showed a much stronger inhibitory potential in ELISA.

### Anti-trypanosomal Activity and Cytotoxicity Analysis

The shortlisted (and validated) compounds were tested in a resazurin survival assay to assess their effects on the growth of *Trypanosoma brucei* cells (**Fig. 7**). Here, suramin - a known drug against HAT [26] - served as positive control. CmpX and risedronate showed trypanocidal effects with EC_50_ (effective concentration leading to 50% cell death) values of 204 nM and 7.7 μM, respectively (**Fig. 7** left panel, bottom and middle). Interestingly, clodronate did not kill the parasites although it could inhibit the *Tb*PEX15–*Tb*PEX6 PPI at nanomolar concentrations in the AlphaScreen, was confirmed in ELISA and is a structural analog of risedronate (left, top panel). Ceftazidime did not influence parasite survival (**Fig. 7** right panel, bottom). Vapreotide and L-selenomethionine affected parasite growth only at the highest concentration of 100 μM (**Fig. 7** right panel, top and middle). Since CmpX and risedronate exhibited potent trypanocidal activity against bloodstream form *T. brucei*, their cytotoxicity was further assessed on human hepatocyte cell line (HepG2) in concentrations up to 100 μM (**Fig. 8A-B**). Blasticidin, a commonly used antibiotic was used as positive control. Both compounds did not exhibit any significant cytotoxicity up to 100 μM.

**Figure 7.**
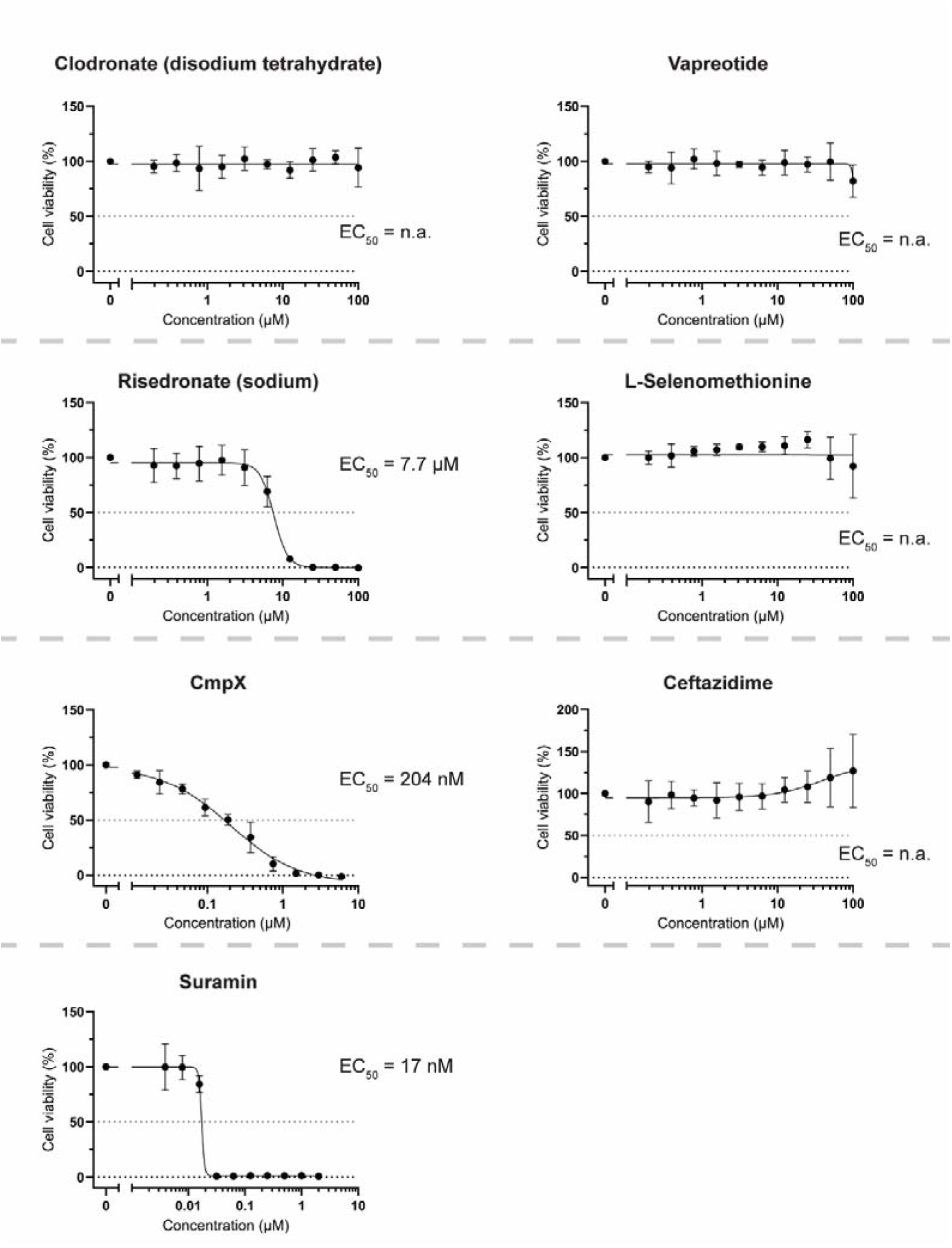
Anti-trypanosomal activities of *Tb*PEX15-PEX6 interaction inhibitors against *T. brucei* bloodstream form parasites. Wildtype *T. brucei* (BF221) cells were treated with different concentrations of inhibitors (two-fold serial dilutions from 100 μM to 195 nM, or 6 μM to 117 nM for CmpX). Risedronate and CmpX have trypanocidal effects with EC_50_ (Effective concentration leading to 50% reduction in parasite viability) values in the low μM or even nM range (7.7 μM and 244 nM, respectively). Clodronate, vapreotide, L-selenomethionine and ceftazidime did affect parasite survival at any concentration. Shown are the results of three biological replicates, error bars indicate ±SD. EC_50_ values were determined via GraphPad Prism 10.0.0.

**Figure 8.**
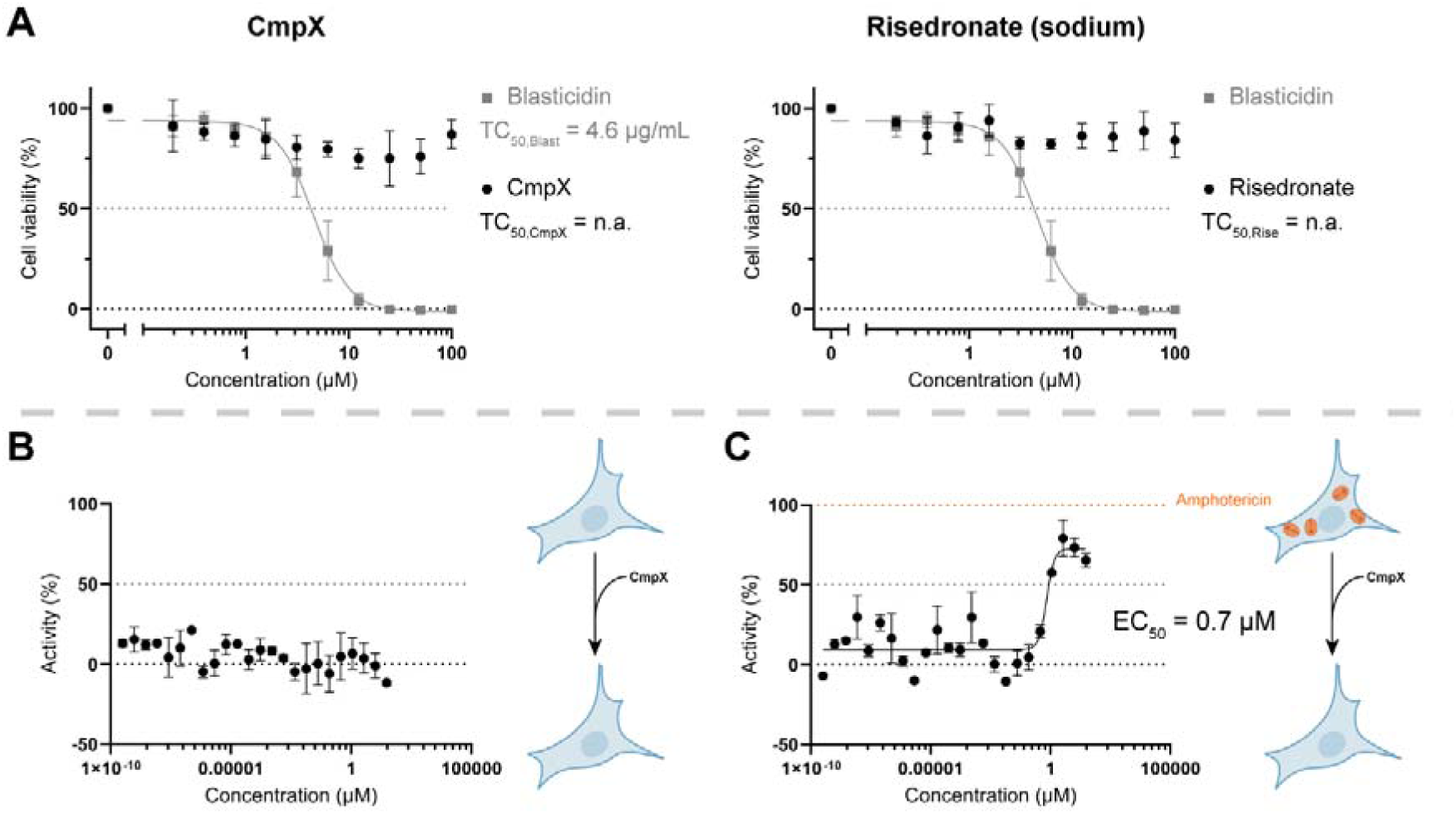
Cytotoxicity and anti-protozoal activity against *T. cruzi* of CmpX and risedronate. (**A**) Cytotoxicity assay of selected compounds CmpX and risedronate on HepG2 cells. Mammalian HepG2 cells were treated with serial dilutions of CmpX and risedronate up to 100 μM. Blasticidin and Hygromycin were used as controls and exhibited EC_50_ values of 4.6 μM and 21.4 μM, respectively. Shown are the results of three biological replicates, error bars indicate ±SD. TC_50_ (Toxic concentrations leading to 50% reduction in viability of mammalian cells) values were determined via GraphPad Prism 10.0.0. (**B**) Cytotoxicity of CmpX on mouse embryonic fibroblasts (NIH/3T3 cell line). (**C**) Anti-protozoal activity of CmpX against *T. cruzi* amastigotes. CmpX was also tested for anti-trypanosomal activity against *T. cruzi*. Mouse embryonic fibroblasts (NIH/3T3 cell line) were infected with *T. cruzi* parasites to establish the intracellular parasite stage called amastigotes. The infected cells were treated with a three-fold serial dilution of CmpX starting from 30.3 μM. Amphotericin B was used as a positive control. CmpX shows potent activity against intracellular *T. cruzi* amastigotes with EC_50_ of 0.7 μM. Shown are the results of three biological replicates, error bars indicate ±SD. EC_50_ values were determined via GraphPad Prism 10.0.0.

### Target Validation of PEX15–PEX6 Inhibitors

The drugs potently inhibited the *Tb*PEX15-PEX6 interaction *in vitro* and exhibited nanomolar trypanocidal activity. To investigate whether the drugs exerted their trypanocidal effect by disrupting the interaction in *T. brucei* parasites, we performed various experiments, including the test for (i) defects in glycosomal protein import by biochemical fractionation using digitonin, (ii) initiation of the RADAR-pathway by analysis of the steady state level analysis of PEX5, (iii) overexpression of *Tb*PEX15 *in cellulo* and (iv) survival assays procyclic parasites in presence and absence of glucose. For the digitonin fractionation, cells were treated with 400 nM CmpX and 10 μM risedronate (concentrations that lead to ∼50% cell death within 48 h of treatment in a viability test, data not shown). The distribution of the cytosolic marker enolase as well as several glycosomal proteins were investigated by immunoblotting (**Fig. S5A**). We did not observe any mislocalization of glycolytic enzymes to the cytosol in presence of compounds compared to H_2_O treatment, indicating that the drugs’ trypanocidal activity might be due to off-target effects. For risedronate, this is in line with its main target being the farnesyl pyrophosphate synthase (FPPS). If the compounds disrupt other interactions or disturb processes with higher affinities than matrix protein import, a mislocalization of glycolytic enzymes is not easily detectable. We next checked for PEX5 levels upon compound treatment. We recently observed PEX5 level reduction upon PEX15 RNAi [24], and hypothesized that a disruption of the PEX15-PEX6 interaction by drug treatment should show a similar effect. However, treatment with CmpX or risedronate did not have any significant effect on PEX5 steady state levels (**Fig. S5B**). For further confirmation, we assessed the compounds’ effect on bloodstream form *T. brucei* in a PEX15 overexpression background. Codon-optimized RNAi-resistant Flag-*Tb*PEX15 overexpression was induced with different concentrations of tetracycline (10 ng/mL and 1 μg/mL, DMSO as control) and the cells were treated with compounds as described for the survival assay above (**Fig. S6**). If the compounds disrupt the PEX15–PEX6 interaction *in cellulo*, overexpression of 2xFlag-*Tb*PEX15 would be expected to shift the EC_50_ to higher values. Suramin and the antibiotic puromycin were used as controls. However, no increase in EC_50_ was observed for the controls, nor the compounds (**Fig. S6B**, top panel). Opposingly, induction of PEX15 overexpression with 1 μg/mL tetracycline caused a slight decrease in EC_50_ for suramin-, puromycin-and risedronate-treated cells (**Fig. S6B**, bottom panel), suggesting that PEX15 overexpression sensitizes BSF *T. brucei* towards the trypanocidal effect of these drugs. This is in line with previous reports of PEX11, another glycosomal membrane protein, overexpression inducing glycosome clustering and affecting cell viability [31,32]. Moreover, both CmpX and risedronate, showed lower EC_50_ values in the PEX15 overexpressing cell line compared to the wild type strain (**Fig. 7**). Since the PEX15 overexpression strain has been maintained under antibiotic selection, it may be more susceptible to drug treatment, or generally less robust. Hence, the observed differences likely represent strain-dependent effects.

To nevertheless confirm the compounds’ inhibitory effect on PEX15 – PEX6 interaction, we also tested the drugs on procyclic form cells in the presence or absence of glucose (**Fig. S7**). Procyclic form (PCF) trypomastigotes mainly metabolize proline for energy supply. In presence of glucose, however, proline-metabolism is downregulated and the cells rely on glycolysis for ATP generation [33]. Disruption of matrix protein import by inhibiting *Tb*PEX15-PEX6 leads to mislocalization of glycolytic enzymes. In the presence of glucose (high-glucose conditions), this causes overproduction of phosphorylated glucose metabolites and ATP depletion ultimately leading to cell death. In the absence of glucose (low-glucose conditions), mislocalization of these enzymes will have no impact on cell survival. Hence, an effect of the compounds on matrix protein import should shift the EC_50_ to lower values in high glucose conditions. Here, blasticidin served as positive control. CmpX and risedronate were tested in concentrations up to 200 μM. CmpX did not affect PCF cell survival independently of glucose availability (**Fig. S7**, top right). Risedronate affected cell survival in both the presence and absence of glucose but seemed to have stronger trypanocidal activity in growth conditions lacking glucose (**Fig. S7**, top left). This again indicates that the compound’s killing effect is most likely not due to targeting of the *Tb*PEX15-*Tb*PEX6 interaction. Based on these results, we conclude that the trypanocidal activity of the identified drugs is mainly due to interference with other essential cellular processes.

### CmpX shows potent and selective activity against *T. cruzi* amastigotes

PEX proteins are highly conserved within trypanosomatid parasites but show very low conservation to human counterparts. To test if the identified drugs are also active against other clinically important trypanosomatid parasites, we extended our study to *Leishmania* and *T. cruzi*. For *Leishmania*, we used *L. tarentolae* promastigotes, a model strain non-pathogenic to humans. Both CmpX and risedronate were tested in concentrations up to 100 μM. Interestingly, no effect of the drugs on cell survival was observed (**Fig. S8**). The positive control – treatment with Amphotericin B – affected cell viability with an EC_50_ of 282.1 nM. Risedronate along with other bisphosphonates are known to inhibit the essential farnesyl pyrophosphate synthase and kill not only *T. brucei*, *T. cruzi* and *L. donovani*, but also *Toxoplasma gondii* and *Plasmodium falciparum* [34–40]. Hence, it was not further investigated regarding its killing effects on *T. cruzi*. Nonetheless, there is no literature reporting CmpX activity against trypanosomatids. Therefore, we tested the drug against *T. cruzi* amastigotes, the intracellular stage of parasites inside infected mammalian host cells. CmpX did not show any cytotoxicity to fibroblast cells alone up to 200 μM tested concentration (**Fig. 8D**). Interestingly, CmpX showed a potent and specific activity against *T. cruzi* amastigotes with an EC_50_ value of 0.7 μM. The reported EC_50_ of currently used drug benznidazole in same assay is >2 μM [41]. The selectivity index (SI), defined as the ratio of mammalian cytotoxicity to trypanocidal activity, reflects both pathogen selectivity and host safety. An SI value of >10 is generally considered favorable in the early stages of drug discovery. In our study, CmpX exhibited an SI of >490 against *T. brucei* and >43 for *T. cruzi*, highlighting its remarkable potency and safety as potential therapeutic agent. All IC_50_, EC_50_ and selectivity index values are summarized in **Table S3 and S4**.

## Discussion

A recent study identified the *Trypanosoma* functional homolog PEX15 within a high-confidence glycosomal membrane protein inventory [24]. PEX15 is essential for *T. brucei* survival, enabling continuous glycosomal matrix protein import through its interaction with PEX6, as demonstrated via Y2H assays. Here, we show that purified recombinant *Tb*PEX15 lacking its C-terminal membrane anchor and full-length *Tb*PEX6 directly interact with each other *in vitro* in a 1:1 ratio, with an apparent K_d_ in the nanomolar range. Due to the low sequence similarity of *Tb*PEX15 to the human functional homolog PEX26, PEX15-PEX6 interaction is an attractive drug target. Along this line, we employed the AlphaScreen technology, which has been successfully used to identify inhibitors of parasite PEX PPIs [1,4]. We established this assay for the *Tb*PEX15-*Tb*PEX6 interaction and screened a drug repurposing library of compounds that are either FDA-approved or in phase II/III of clinical trials and thus have passed toxicity evaluations. After several quality controls, six compounds were identified that selectively disrupt the *Tb*PEX15–*Tb*PEX6 interaction *in vitro*. Two of these compounds, CmpX and risedronate, exhibited potent and selective trypanocidal activity against *T. brucei*; however, subsequent *in cellulo* assays showed no significant impact on glycosome biogenesis. A block of the PEX15 – PEX6 interaction is expected to interfere with the recycling of the import receptor PEX5. In this scenario, the ubiquitinated receptor would accumulate at the peroxisomal membrane, which may trigger pexophagy or degradation of the receptor in the RADAR (Receptor Accumulation and Degradation in the Absence of Recycling) pathway [42]. However, such a decrease in steady-state concentration of PEX5 is not detected. This suggests that the drugs’ trypanocidal activity might arise from interference with other essential cellular processes rather than from the disruption of glycosome biogenesis. Nonetheless, CmpX showed potent trypanocidal activity against clinically important *T. cruzi* intracellular amastigotes, with no cytotoxicity against the infected host cell. In this aspect, CmpX shows even better *in vitro* activity against amastigote stage parasites (EC_50_ 0.7 µM) than the currently used drug benznidazole (EC_50_ >2 µM) in a similar assay [43].

### Bisphosphonates

The bisphosphonates risedronate and clodronate disrupted the *Tb*PEX15 – PEX6 interaction in both AlphaScreen assay and ELISA. However, structurally similar bisphosphonates, specifically zoledronic acid, etidronic acid, minoronic acid, pamidronate, alendronic acid and ibandronate had no specific inhibitory effect. Despite a clear toxicity for trypanosomatids, target validation studies revealed that glycosome-related function, including protein import into the organelle, are not affected. The lack of interference with glycosomal functions indicates that the drugs’ trypanocidal activity might be due to off-target effects.

Bisphosphonates are primarily known for their use in treating various skeletal disorders [50]. They are chemically stable structural analogues of inorganic pyrophosphate (PPi) and as such have a very high affinity for bone minerals as they bind to hydroxyapatite crystals. Along this line, they likely inhibit bone calcification and hydroxyapatite breakdown [51,52]. Clodronate is an early non-nitrogen containing bisphosphonates that is thought to be incorporated into newly formed ATP, thereby rendering it nonhydrolyzable. This ATP analog is thus cytotoxic to osteoclasts. Risedronate on the other hand belongs to the nitrogen-containing bisphosphonates that are not incorporated in ATP but rather inhibit farnesyl pyrophosphate synthase (FPPS) [53,54]. This prevents post-translational modification, specifically isoprenylation, of proteins, which ultimately leads to osteoclast apoptosis [55]. Nitrogen-containing bisphosphonates had already been shown to have killing effects on several protozoan organisms including *T. brucei*, *T. cruzi* and *L. donovani* [36]. It was shown that bisphosphonates inhibit *T. cruzi* FPPS [38] and in *T. brucei* it was found that FPPS is an essential protein and growth can be inhibited by bisphosphonates [34]. These observations explain the trypanocidal effect of risedronate observed within this study.

There is increasing interest in developing human PEX protein interactions for anti-cancer treatments [21,22,56]. Particularly, silencing of PEX26, the PEX15 counterpart in humans, kills drug resistant cancer cells and is thus proposed as an unconventional mode for therapy [22]. Therefore, the established screen for *Trypanosoma* PEX15-PEX6 may be adapted to human PEX26-PEX6 to identify of inhibitors for anticancer therapy development.

## Conclusion

This study provides the basis for further dissection of the *Trypanosoma* PEX15-PEX6 interaction and platform for the identification of inhibitors of this interaction. Since *T. brucei* infections are now limited, the inhibitor screening procedure could be easily adapted for related parasites, i.e., *T. cruzi* and *Leishmania,* which still lead to significant infections and mortality, hence require new efficient therapies.

## Author contribution

LH, VK, RE conceived and planned the experiments. LH, BT and FT performed all experiments. VK, RE supervised the work. LH wrote the manuscript with support of VK and RE.

## Acknowledgements

This work was supported by the Deutsche Forschungsgemeinschaft (ER 178/17-1) to RE, and InnovationsFoRUM grants of the Ruhr-University Bochum IF-009N-22, IF-018N-22) to RE. Authors thank Prof. Dr. Paul Michels for kindly providing various *Trypanosoma* antibodies.

## Methods

### Cloning

*Trypanosoma* expression plasmid constructs and cloning strategies are listed in **Table S3**, and oligonucleotide sequences are listed in **Table S4**. Sequences of the constructs were verified for all constructs by automated Sanger sequencing.

### Protein expression

His_6_-*Tb*PEX15ΔTM (1-320aa) (from pET28a plasmid with TEV cleavage site) was expressed in *E. coli* BL21 pRIL (*DE3*) at 18°C overnight after induction with 0.5 mM IPTG. GST-*Tb*PEX6 (from pGEX-4T-2 plasmid) full-length was expressed in *E. coli* BL21 Star (*DE3*) at 20°C for 20 h. Overexpression of GST-His_6_ protein (from pET42b+) was performed at 30°C for 4 h in the presence of 1 mM IPTG. Cells were harvested at 4,500 × *g* (SLA-3000) for 5 min and the pellet stored at-70°C.

### Protein purification

#### His_6_-TbPEX15ΔTM and GST-TbPEX6

Cells were thawed on ice and according to the pellet’s weight, ten times volume of lysis buffer (50 mM Tris, 150 mM NaCl, pH 7.9, supplemented with 5 µg/mL Antipain, 2 µg/mL Aprotinin, 0.35 µg/mL Bestatin, 6 µg/mL Chymostatin, 2.5 µg/mL Leupeptin, 1 µg/mL Pepstatin A, 2.5 µg/mL DNAse, 1 mM PMSF, and 1 mM DTT) was added and the pellet resuspended. For homogenization, the pellet was dounced four times and then lysed using EmulsiFlex (Avestin). The homogenate was centrifuged at 4,500 × *g* (SLA-3000, Thermo Scientific) for 15 min at 4°C. The pellet contains non-lysed cells. The lysate containing supernatant was centrifuged at 14,000 × *g* (SS-34 rotor, Sorvall® Thermo Scientific) for 1 h at 4°C for removal of cell debris. The supernatant was taken forward and loaded onto a 5 mL Ni-NTA column (Protino®, Macherey-Nagel) for His_6_-tagged *Tb*PEX15 protein or 1 mL GST/4B column (Protino®, Macherey-Nagel) for GST-tagged *Tb*PEX6. After loading, the column was washed with washing buffer (50 mM Tris, 150 mM NaCl, pH 7.9). After 10 column volumes (CVs) washing, gradient elution from 0 to 100% with elution buffer I (50 mM Tris, 150 mM NaCl, pH 7.9, 300 mM imidazole) or elution buffer II (50 mM Tris, 150 mM NaCl, pH 7.9, 20 mM glutathione) was performed. The target protein-containing fractions (determined by SDS-PAGE) were pooled and concentrated using an Amicon® Ultra Filter tube with 10 kDa molecular weight cut-off (MWCO) or 30 kDa MWCO, respectively. The concentrate was then further purified and buffer-exchanged by a Superdex^®^ 200 HiLoad 26/600 prep grade or Superdex^®^ 200 increase 10/300 GL (GE Healthcare, now Cytiva) size-exclusion chromatography using HEPES buffer (25 mM HEPES, 150 mM NaCl, pH 7.5). For generation of tag-free *Tb*PEX15 protein, pooled fractions after Ni affinity chromatography were treated with TEV protease at 4°C overnight without agitation. Reverse nickel affinity chromatography with wash buffer and elution buffer I was performed using a 5 mL Ni-NTA column (Protino®, Macherey-Nagel) for removal of TEV-protease followed by size exclusion chromatography using HEPES buffer. Pure protein fractions were pooled and again concentrated. All protein purification steps were performed at 4°C.

#### GST-His_6_ protein

Cells were thawed on ice and resuspended in ten times volume of lysis buffer (1x PBS, pH 7.4, supplemented with 5 µg/mL Antipain, 2 µg/mL Aprotinin, 0.35 µg/mL Bestatin, 6 µg/mL Chymostatin, 2.5 µg/mL Leupeptin, 1 µg/mL Pepstatin A, 2.5 µg/mL DNAse, 1 mM PMSF, and 1 mM DTT) according to the pellet’s weight. For homogenization, the sediment was dounced four times and then lysed using EmulsiFlex (Avestin). The homogenate was centrifuged at 4,500 *x g* (SLA-3000, Thermo Scientific) for 15 min at 4°C. The supernatant was further centrifuged at 14,000 *x g* (SS-34 rotor, Sorvall® Thermo Scientific) for 1 h at 4°C. The protein was purified in batch using Protino® Glutathione Agarose 4B beads. The protein was eluted using 1 mM to 10 mM reduced glutathione. Eluate fractions were pooled and dialyzed in 1x PBS, pH 7.4. All protein purification steps were performed at 4°C.

Protein concentrations were determined by Bradford assay (Coomassie Plus assay kit, Thermo Scientific). Aliquots were snap-frozen in liquid nitrogen and stored at-70°C until further use.

#### *In vitro* pull-down assay

The *in vitro* pull-down assay was performed at 4°C. GST-*Tb*PEX6 (10 μM) was incubated with equilibrated Glutathione agarose 4B beads (Protino^®^, Macherey Nagel) in wash buffer (50 mM Tris, 150 mM NaCl, pH 7.9) with light agitation. After an incubation for 1 h, the flow-through was collected by centrifugation. The beads were washed three to five times with wash buffer. His_6_-*Tb*PEX15ΔTM was added to the beads and incubated for 1 h further, after which the flow-through was collected by centrifugation. The beads were washed again three to five times with wash buffer. Subsequently, the proteins were eluted twice with elution buffer (wash buffer supplemented with 20 mM reduced glutathione). Lastly, the beads were boiled at 95°C after addition of SDS-Laemmli buffer. The collected fractions were analyzed by SDS-PAGE. For analysis of binding stoichiometry, the protein-bound beads (GST-*Tb*PEX6 + His_6_-*Tb*PEX15ΔTM) were incubated with GST-PreScission protease for 2 h at 4°C. The flow-through was collected and subjected to size exclusion chromatography (SEC). The beads were further washed three times with wash buffer, after which the remaining proteins were eluted with reduced glutathione as described above.

### Size exclusion chromatography

To investigate binding stoichiometry, gel filtration analysis was performed with an ÄKTApurifier system (GE Healthcare, Freiburg, Germany) on a Superose™6 PC 3.2/30 column (GE Healthcare, Freiburg, Germany) equilibrated with 50 mM Tris, 150 mM NaCl, pH 7.9. 50 μL of recombinant proteins or protein mixtures was loaded and subsequently, 50 μL fractions were collected.

The column was calibrated with thyroglobulin (669 kDa), ferritin (440 kDa), aldolase (158 kDa), ovalbumin (44 kDa), carbonic anhydrase (29 kDa), ribonuclease A (13.7 kDa) and aprotinin (6.5 kDa).

### AlphaScreen – Cross Titration

For assay establishment, the recombinant proteins His_6_-*Tb*PEX15ΔTM and GST-*Tb*PEX6 were cross-titrated. All steps were performed at room temperature (RT). A serial titration from 0 - 300 nM of both proteins was performed. 5 µL of GST-*Tb*PEX6 were incubated with 5 µL of His_6_-*Tb*PEX15 and 5 µL of buffer for 30 min. 5 µL of Ni-NTA acceptor beads (Revvity, 1:250 dilution in assay buffer) were added for binding to His_6_-*Tb*PEX15ΔTM in the dark for 15 min. Subsequently, 5 µL of donor beads (Revvity, 1:250 dilution in assay buffer) were added to bind GST-*Tb*PEX6 proteins for another 40 min in the dark. The Alpha signal was monitored with a Cytation 5 plate reader (BioTek®) with the gain value set at 180.

### AlphaScreen – Saturation assay

As described for the cross-titration above. 1, 3 and 10 nM GST-*Tb*PEX6 were incubated with a serial dilution His_6_-*Tb*PEX15ΔTM starting from 300 μM.

### AlphaScreen – Displacement assay

10 or 20 nM His_6_-*Tb*PEX15ΔTM were incubated with 1 or 2 nM GST-*Tb*PEX6. After a 30 min incubation at RT, a serial dilution of tag-free *Tb*PEX15ΔTM was added to the protein mix and incubated for 1 h after which the beads were added as described for cross titration above.

### AlphaScreen - Primary Screen

The AlphaScreen assay was performed in light grey, low binding, 384 well AlphaPlate (Revvity). Protein aliquots were thawed on ice and then diluted in assay buffer (HEPES buffer with 0.5% BSA, 0.05% Tween-80). Drug plates were thawed at RT and centrifuged at 2,000 rpm (SX4750-A, Beckman Coulter) for 2 min. His_6_-*Tb*PEX15ΔTM and GST-*Tb*PEX6 were incubated together in a 1:1 ratio at RT for 30 min with a concentration of each 25 nM. Then 10 μL of the protein solution were incubated with 5 μL of 50 μM/ 10 μM/ 15 μg/mL (depending on plate) drugs for 1 h in a 384-well plate, after which 5 μL Ni-NTA acceptor beads were added. GSH donor beads were added after 15 min incubation. The proteins were at a final concentration of 10 nM, drugs at 10 μM/ 2 μM/ 3 μg/mL and beads each at 0.8 μg/mL in a final reaction volume of 25 μL. Alpha signal was detected after 40 min with Cytation 5 plate reader (BioTek®) with the gain value set at 180. As positive control, buffer supplemented with the respective solvent in which the drugs were diluted initially was added instead of drugs (highest signal). Beads only served as negative control (lowest signal). Addition of NaCl at a final concentration of 1 M in 25 μL showed that the interaction can be disrupted and hence, served as another control.

### AlphaScreen - Counter Screen

Drugs that showed > 50% inhibition or an RZ-score <-3 in the primary screen were tested in a counter screen against GST-His_6_ as well as His_6_-*Tb*PEX15ΔTM and GST-*Tb*PEX6 to validate that the observed disruption is specific for the *Tb*PEX15-*Tb*PEX6 interaction. The assay was performed as described above for the primary screen.

### Dose-response AlphaScreen

Drugs that showed > 80% inhibition of *Tb*PEX15-*Tb*PEX6 interaction and < 20% inhibition of GST-His_6_ were procured as 10 mM stocks from MedChemExpress. Dose-response measurements were performed with three technical replicates. A solution of His_6_-*Tb*PEX15ΔTM and GST-*Tb*PEX6 was incubated for 30 min at RT. The protein solution was then incubated at a concentration of each 12.5 nM with compound in assay buffer (HEPES buffer + 0.5% BSA, 0.05% Tween-80) for 1 h. Acceptor and donor beads were added as described above. Final compound concentrations ranged from 50 nM to 100 μM. Half maximal inhibitory concentrations (IC_50_) were determined from three biological replicates using GraphPad Prism 10.0.0. For calculation, the equation: log(inhibitor) vs. response -- Variable slope (four parameters) was used.

### ELISA assay

Shortlisted compounds were validated in an orthogonal ELISA assay, which was performed in 96 well microtiter plates (Immulon™ 2 HB 96-Well Microtiter EIA Plate, ImmunoChemistry Technologies) at RT. 1 µg His_6_-*Tb*PEX15ΔTM was coated in wells at 4°C overnight. Wells were washed twice with 250 μL ELISA buffer (50 mM Tris, 137 mM NaCl, pH 8.0 supplemented with 0.1% Tween-20 to remove unbound protein. After blocking with 200 μL blocking buffer (3% BSA in ELISA buffer) for 1 h at RT and another wash step, 100 μL of serial dilution of compounds was incubated with coated *Tb*PEX15 and the unbound proteins were washed out three times with PBS. To these wells, 100 µl of GST-*Tb*PEX6 was added to reach final concentration of 10 μM and incubated for 30 min further. After three washes with ELISA buffer, bound GST-*Tb*PEX6 was detected with mouse monoclonal HRP-coupled anti-GST antibody (Sigma-Aldrich, 1:2,000 v/v in ELISA buffer) for 1 h at RT. Following four washes, substrate 3,3′,5,5′-tetramethylbenzidine (TMB, Thermo Fisher Scientific) was added. The colorimetric reaction was terminated after 20 min by adding H_2_SO_4_, and the absorbance was measured at wavelength of 450 nm.

### Culture maintenance

*Trypanosoma brucei* BSF strain Lister 427 (termed hereafter as BSF427) cells were maintained in HMI-11 medium [57] in logarithmic phase (below 2 × 10^6^ cells/ml as described in [58]) in a humidified incubator at 37°C and 5% CO_2_. Procyclic form (PCF) trypanosomes (strain 29-13) were maintained in SDM79 medium [59] without NaHCO_3_ at 28°C without CO_2_ in logarithmic phase (below 1-2 × 10^6^ cells/ml as described in [60,61]). *Leishmania tarentolae* promastigotes (LEXSY host P10, Jena Bioscience) were grown in LEXSY Broth BHI medium (Jena Biosciences) supplemented with 7.5 μg/mL hemin. HepG2 (hepatocyte) cells were used to study the cytotoxicity of compounds. Cells were maintained at 37°C with 5% CO_2_ in high-glucose Dulbecco’s Modified Eagle’s Medium (DMEM) supplemented with L-glutamine (4 mM). All media were used with 10% heat-inactivated fetal bovine serum (FBS), penicillin (100 units/L) and streptomycin (100 mg/L).

### Resazurin Survival Assay in blood-stream form *T. brucei*

*T. brucei* BSF427 cells or BSF 90-13 cells transfected with an overexpression construct for codon-optimized *Tb*PEX15 in HMI-11 medium [57] were seeded at a density of 4×10^3^ cells/mL (100 μL per well) in 88 wells of a clear, polystyrene 96-well plate. Medium only was added to 8 wells as control. Compounds at final concentrations of 195 nM to 100 μM (2-fold serial dilution) of L-selenomethionine, ceftazidime, clodronate, risedronate and vapreotide, 12 nM to 6 μM for CmpX or 4 nM to 2 μM for suramin (control) were tested in four technical replicates. The final volume was 200 μL. After 66 h incubation at 37°C and 5% CO_2_ in humidified atmosphere, 25 μL resazurin solution (0.1 mg/mL) in HBSS was added to all wells. Cell viabilities were measured quantitatively by means of fluorescence detection after 6 h incubation using TECAN Infinite^®^ M Nano^+^ at 530/585 nm and 530/570 nm for background correction. Half maximally effective concentrations (EC_50_) were determined from three biological replicates using GraphPad Prism 10.0.0. For calculations, the equation: log(inhibitor) vs. response -- Variable slope (four parameters) was used.

### Resazurin Survival Assay in procyclic form *T. brucei*

Two-fold serial dilutions of CmpX and risedronate ranging from 781 nM to 200 μM (8 wells in each row) were prepared in clear, polystyrene 96-well plates. One row was filled with 100 μL medium. 100 μL of *T. brucei* procyclic form cells (1×10^6^ cells/mL in SDM79 medium) were added to the wells with 100 μL of prepared compounds or medium. Another row with 200 μL medium were included as control. The outermost wells were also filled with 200 μL medium to reduce evaporation during the following incubation in a non-humidified atmosphere. The cells were incubated at 28°C for a total of 72 h. 25 μL of resazurin reagent (0.1 mg/mL in HBSS) was added to all wells after 70 h incubation and further incubated for 2 h at 28°C. To compare the effect of the compounds in high-glucose vs. low-glucose conditions, SDM79 medium was either used with 10 mM glucose (high-glucose condition) or in the absence of any glucose but with additional 50 mM N-acetyl glucosamine to hinder uptake of residual glucose from the FBS used in the medium.

### Resazurin Survival Assay in promastigote form *L. tarentolae*

As for PCF trypanosomes, only the inner 58 wells of clear, polystyrene 96-well plates were using for the cell viability assay while the outer wells were filled with 200 μL medium to reduce evaporation. Accordingly, two-fold serial dilutions of CmpX and risedronate ranging from 391 nM to 100 μM (8 wells in each row). One row was filled with 100 μL medium. 100 μL of *L. tarentolae* cells (5×10^4^ cells/mL in LEXSY Broth BHI medium) were added to these wells. One row with 200 μL medium only was prepared and included as control. The cells were incubated at 28°C for 72 h. 25 μL of resazurin reagent (0.1 mg/mL in HBSS) was added to all wells following the 67-h incubation and further incubated for 5 h at 28°C.

### Resazurin Survival Assay in human HepG2 cells

The cytotoxicity assay was performed in 96-well plates using a resazurin-based cell viability assay. [62] Cells were seeded in wells at a density of 5×10^3^ cells per well and incubated at 37°C overnight. Two-fold serial dilutions of compounds were prepared ranging from 195 nM to 100 µM (10 dilutions in total) culture medium. After removing the medium from the overnight cultured cells, compound dilutions were added to each well and incubated for 72 h. After 68 h of incubation, 25 μL resazurin solution (0.1 mg/mL) in HBSS was added to all wells for 4 h incubation. Cell viabilities were measured quantitatively by means of fluorescence detection using TECAN Infinite^®^ M Nano^+^ at 530/585 nm and 530/570 nm for background correction. Results were collected from at least three biological replicates with four technical replicates in each. Control (with 0 µM of compounds) was set to 100% cell viability and used for normalization. Data was analyzed using curve fitting with a dose-response model (EC_50_ shift, X is concentration) in GraphPad Prism 10.0.0 to calculate TC_50_ values. For calculation, the equation: log(inhibitor) vs. response -- Variable slope (four parameters) was used.

### Digitonin fractionation

BSF427 cells were treated with 400 nM CmpX, 10 μM risedronate or H_2_O for 48 h. Biochemical digitonin fractionation was performed according to the following protocol adapted from [63]. Cells were harvested at 1,500 × *g*. The harvested samples were resuspended in 250 mM sucrose and 2.5 µg/mL Leupeptin in PBS (phosphate-buffered saline, pH 7.4). Protein concentration was estimated using the Bradford method. Following the protein estimation, samples with protein corresponding to ∼7 μg were treated with increasing amounts of digitonin from 0.01 mg to 1 mg of digitonin/mg of protein (diluted in PBS with 250 mM sucrose). For the positive control, cells were treated with 1% Triton-X 100 representing the complete release of all proteins by dissolving all membranes. After adding digitonin, the suspension was vortexed with medium intensity for 3 sec and incubated at 37°C for 2 min with shaking (600 rpm, Thermo mixer). The incubated samples were centrifuged at 16,100 × *g* for 30 min at 4°C. The resulting supernatant was further analyzed by immunoblotting using antibodies for enolase (cytosolic marker) and aldolase (glycosomal matrix marker).

## Supplementary Information

**Table S1:** IC_50_ and EC_50_ values of validated hits from drug repurposing library. (All values in **μ**M)

| Compound | IC<sub>50</sub>,<br>AlphaScreen | EC<sub>50</sub>,<br>*T. brucei*<br>(BSF) | EC<sub>50</sub>,<br>*T. brucei*<br>(PCF) | EC<sub>50</sub>,<br>*L.*<br>*tarentolae* | EC<sub>50</sub>,<br>*T. cruzi* | TC<sub>50</sub>,<br>HepG2 |
| --- | --- | --- | --- | --- | --- | --- |
| Ceftazidime | > 100 | > 100 | n.a. | n.a. | n.a. | n.a. |
| Clodronate | 0.199 | > 100 | n.a. | n.a. | n.a. | n.a. |
| CmpX | 0.102 | 0.204 | > 200 | > 100 | 0.7 | > 100 |
| L-Seleno-methionine | 3.06 | > 100 | n.a. | n.a. | n.a. | n.a. |
| Risedronate | Not calculable | 7.7 | > 200 | > 100 | n.a. | > 100 |
| Vapreotide | 18.73 | > 100 | n.a. | n.a. | n.a. | n.a. |

*n.a.* not applicable

**Table S2:** Selectivity indices.

| Compound | Selectivity index<br>*T. brucei* | Selectivity index<br>*T. cruzi* |
| --- | --- | --- |
| Ceftazidime | n.a. | n.a. |
| Clodronate | n.a. | n.a. |
| CmpX | > 490 | > 43 |
| L-Seleno-methionine | n.a. | n.a. |
| Risedronate | > 12 | n.a. |
| Vapreotide | n.a. | n.a. |

**Table S3:**
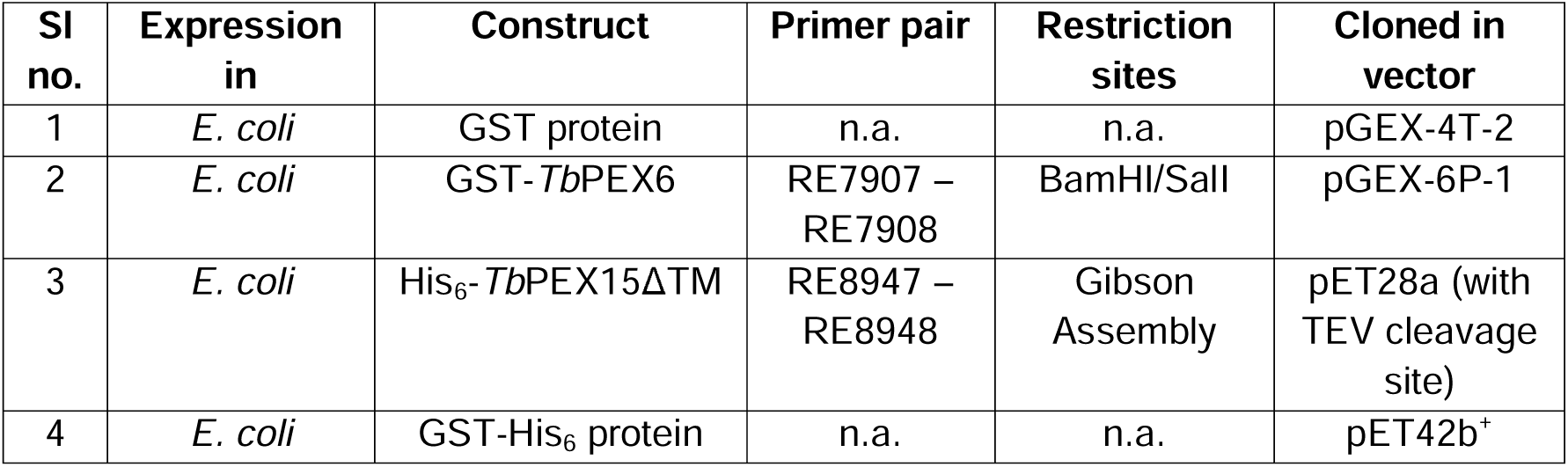
Strains and Plasmids.

**Table S4:**
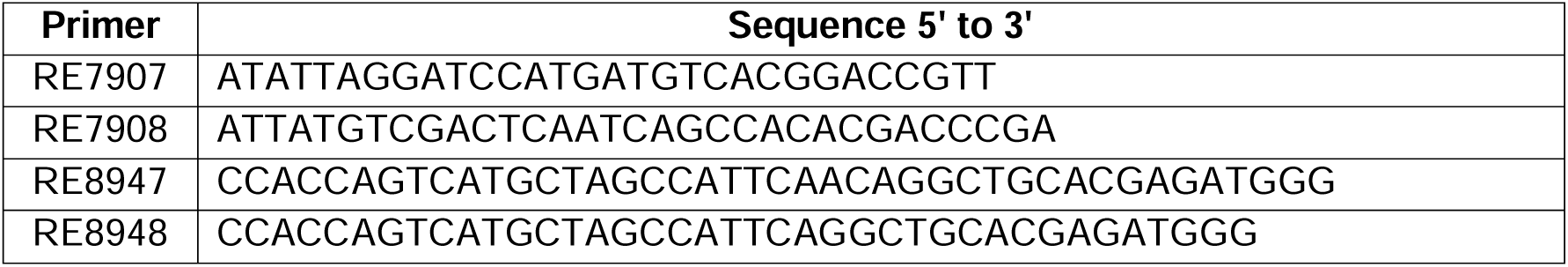
Oligonucleotides.

## Supplementary Figures

**Figure S1.**
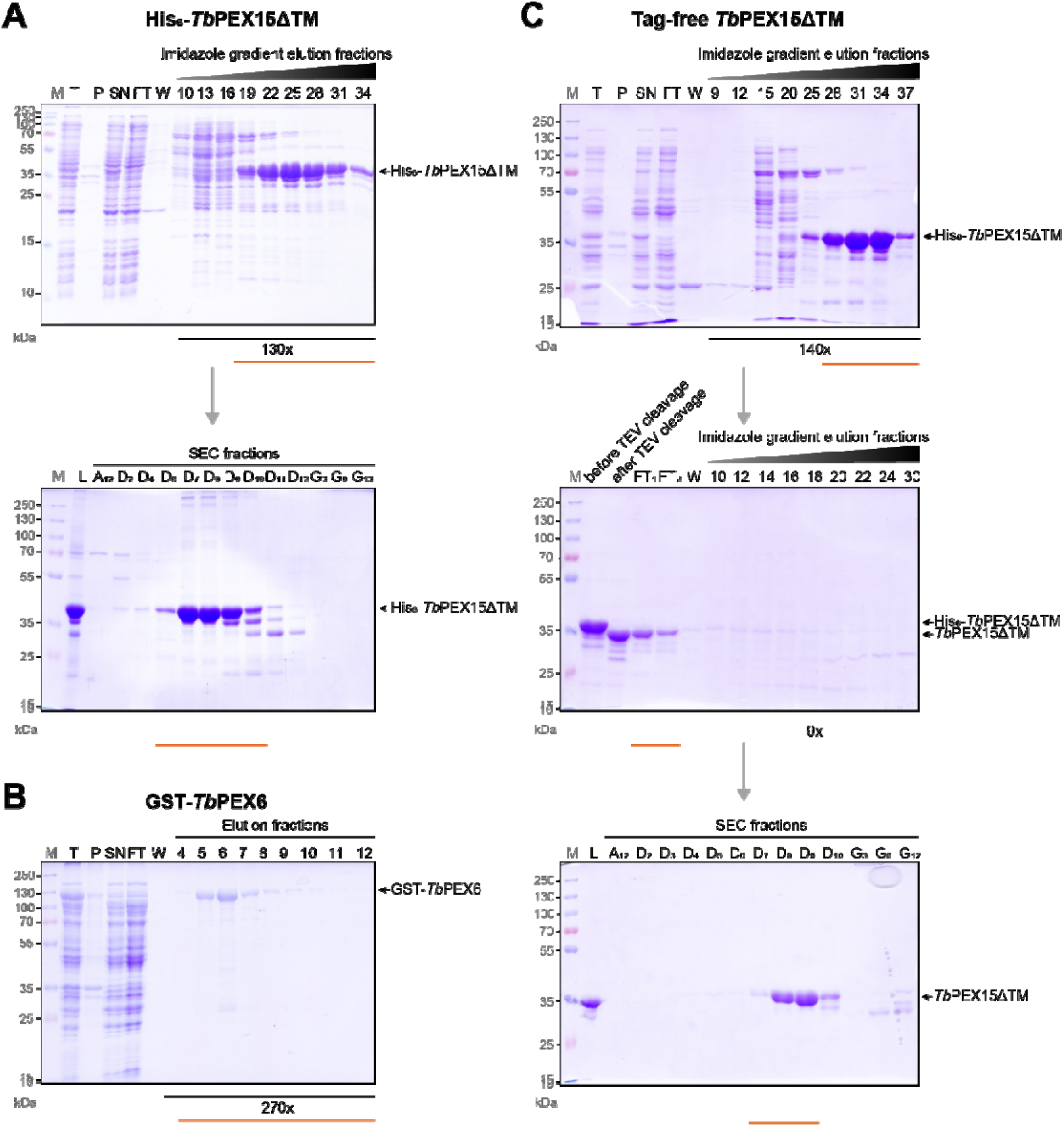
**Purification of recombinant His_6_-tagged and untagged *Tb*PEX15**Δ**TM (1-320aa) and GST-*Tb*PEX6 (*full-length*).** After overexpression, the proteins were purified according to the method described. Affinity chromatography and size exclusion chromatography was performed using the ÄKTA start system. **(A)** His_6_-*Tb*PEX15ΔTM (1-320aa) was affinity purified using a HisTrap column. Bound protein was eluted with a gradient of up to 300 mM imidazole and fractions were analyzed by SDS-PAGE. His_6_-*Tb*PEX15ΔTM with a molecular weight of 36 kDa could be recovered from fractions 22-34, which were pooled and concentrated. **(B)** Size exclusion chromatography allowed removal of further contaminations. GST-*Tb*PEX6 was purified using Protino^®^ GSH agarose column. Isocratic elution yielded pure protein. Elution fractions were pooled and concentrated. **(C)** For generation of untagged *Tb*PEX15ΔTM (1-320aa), His_6_-*Tb*PEX15ΔTM (1-320aa) was affinity purified, followed by tag-cleavage utilizing TEV protease. After cleavage, the protein solution was again submitted to Ni affinity chromatography. The PEX15 protein containing flow-through was further purified by size exclusion chromatography. Pooled fractions are indicated by the orange bar.

**Figure S2.**
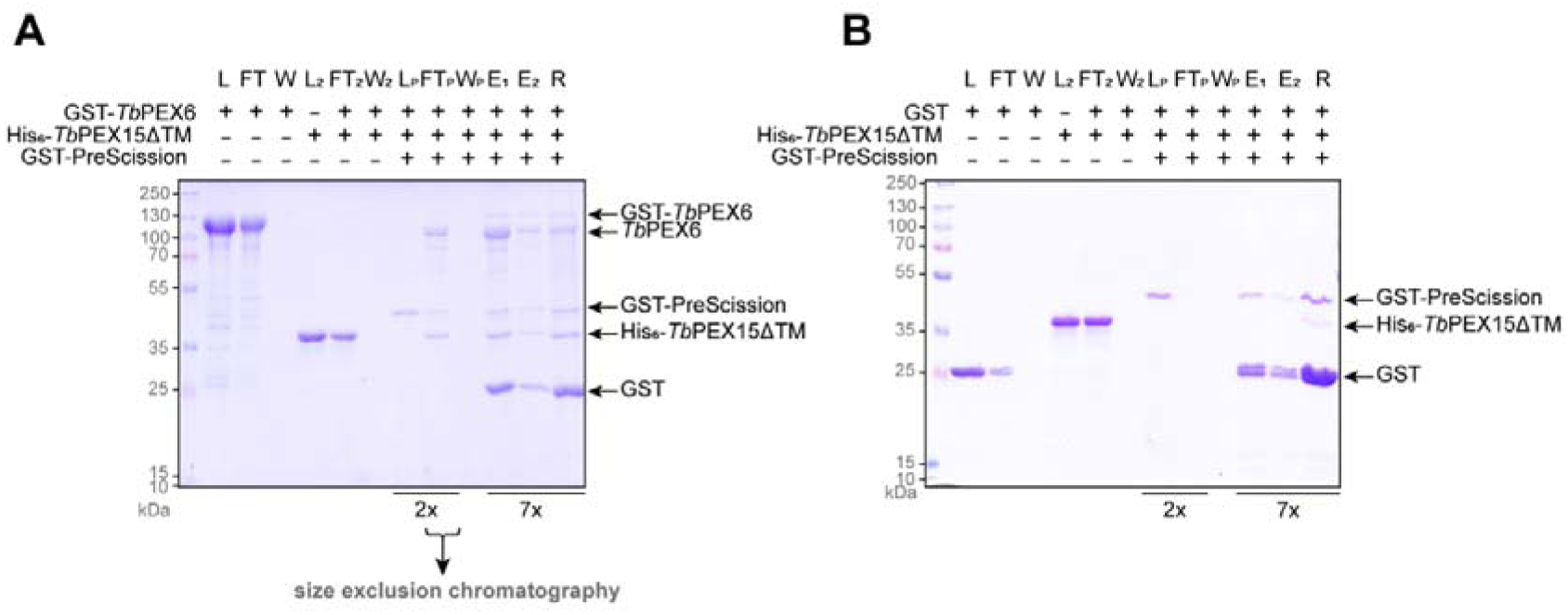
*In vitro* pull-down assay as described in **Fig. 1** with the exception that prior to glutathione elution, protein-bound beads were incubated with PreScission protease for 2 h at 4°C. The flow-through (FT_P_) is collected and subjected to size exclusion chromatography via Superose™6 PC 3.2/30 column (bottom panel).

**Figure S3.**
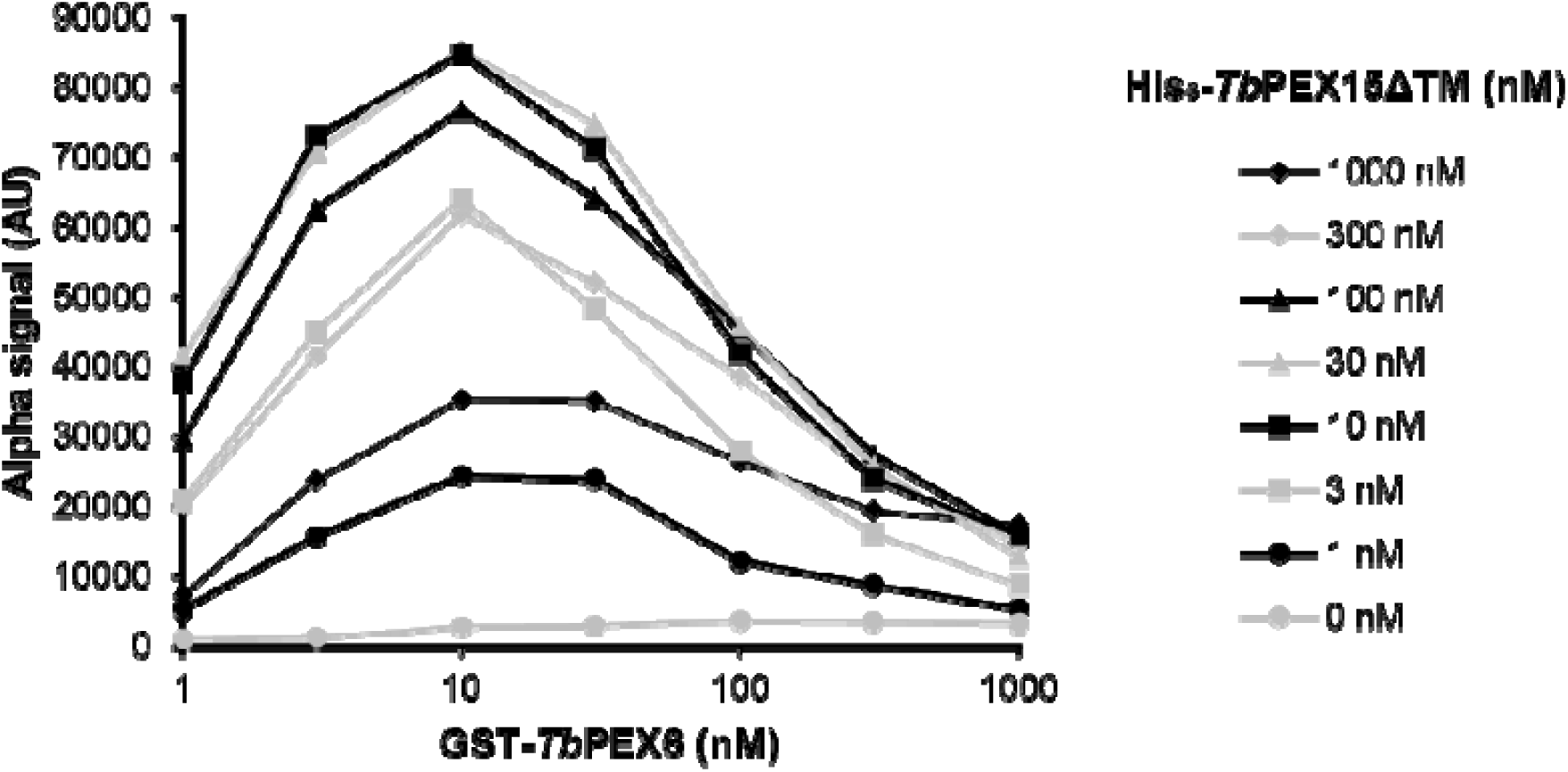
Cross titration of *Tb*PEX15 and *Tb*PEX6 in AlphaScreen. The Alpha assay was performed with 20 ug/ml GSH-donor and Ni-Chelate acceptor beads in 25 mM HEPES, 150 mM NaCl, pH 7.5, supplemented with 0.5% BSA and 0.05% Tween-80 in a reaction volume of 25 ul. Recombinant full-length GST-*Tb*PEX6 and His_6_-*Tb*PEX15ΔTM were cross-titrated with following concentrations: 0, 1, 3, 10, 30, 100, 300, 1000 nM. The Alpha signal in arbitrary units (AU) is plotted against the GST-*Tb*PEX6 concentration in nM. His_6_-*Tb*PEX15ΔTM concentrations are plotted as: 0 nM (grey, dots), 1 nM (black, dots), 3 nM (grey, squares), 10 nM (black, squares), 30 nM (grey, triangles), 100 nM (black, triangles), 300 nM (grey, rhombi), 1000 nM (black rhombi). The maximal Alpha signal is observed at a concentration of 10 nM for each protein. Above these concentrations, the signal drops due to the hooking effect (described in methods section).

**Figure S4.**
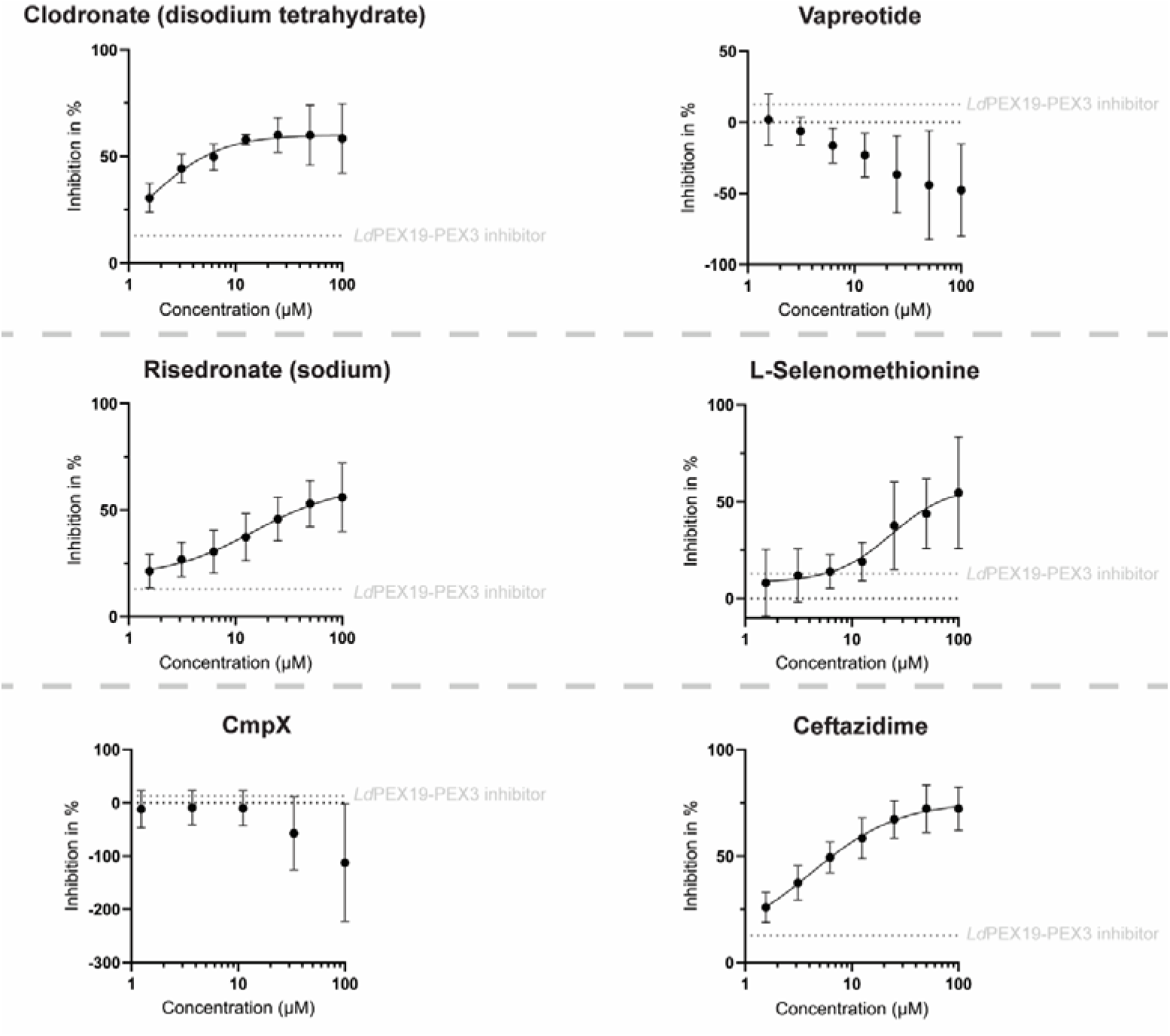
Hit Validation by ELISA. The shortlisted compounds clodronate, vapreotide, risedronate, L-selenomethionine, CmpX and ceftazidime were tested in an enzyme-linked immunosorbent assay (ELISA) in seven concentrations from a two-fold dilution series starting from 100 μM for disruption of the *Tb*PEX15-*Tb*PEX6 interaction. 1ug His_6_-*Tb*PEX15ΔTM was coated in wells of a 96 well plate, blocked with 5% fat-free milk in Tris-buffered saline followed by binding of GST-*Tb*PEX6 in presence of the inhibitors. For CmpX, a three-fold dilution series was performed. The data indicated dose-dependent inhibition for clodronate, vapreotide, risedronate and L-selenomethionine. After a total incubation time of 90 min, an HRP-coupled αGST antibody was used for recognition of bound PEX6, followed by 3,3′,5,5′-tetramethylbenzidine (TMB)-based signal detection. CmpX and vapreotide interfered with signal detection and increasing compound concentrations lead to increasing ELISA signal. As negative control, an inhibitor that was identified from the same library as a *Ld*PEX19-PEX3 inhibitor was used (indicated by the grey dotted line).

**Figure S5.**
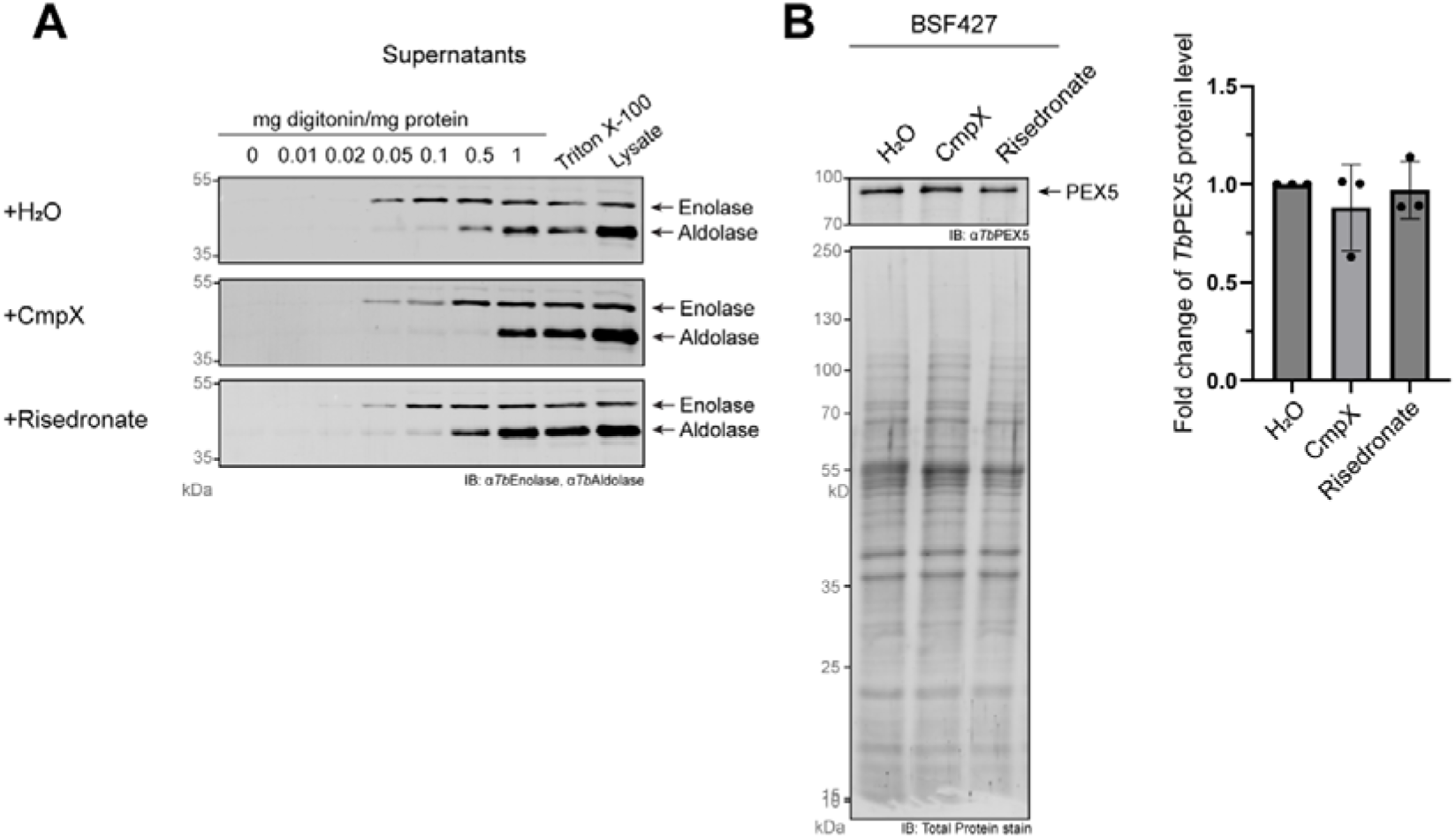
Target validation of CmpX and risedronate in *T. brucei* parasites. (**A**) Biochemical fractionation utilizing digitonin was performed to investigate the effect of the compounds on glycosomal matrix protein import. Disruption of PEX15-PEX6 interaction is expected to result in a phenotype like the one observed for PEX15 and PEX6 knock-down showing partial mislocalization of glycosomal proteins. *T. brucei* bloodstream form cells were treated with 400 nM CmpX, 10 μM risedronate or equivalent amount H_2_O as control for 48 h. Following treatment, cells were harvested and subjected to treatment with different concentrations of digitonin. Triton-X100 served as a positive control, releasing all proteins by dissolving all membranes. The supernatants were analyzed by immunoblotting with antibodies against glycosomal matrix marker aldolase and enolase as a cytosolic marker as well as a loading control. (**B**) Lysates of treated cells were assessed for their steady state PEX5 levels. Samples were analyzed by immunoblotting, a representative blot is shown in the left panel, where enolase was used as loading control. PEX5 bands were normalized to total protein stain and fold change was calculated. Treatment with CmpX or risedronate does not have significant effect on PEX5 levels.

**Figure S6.**
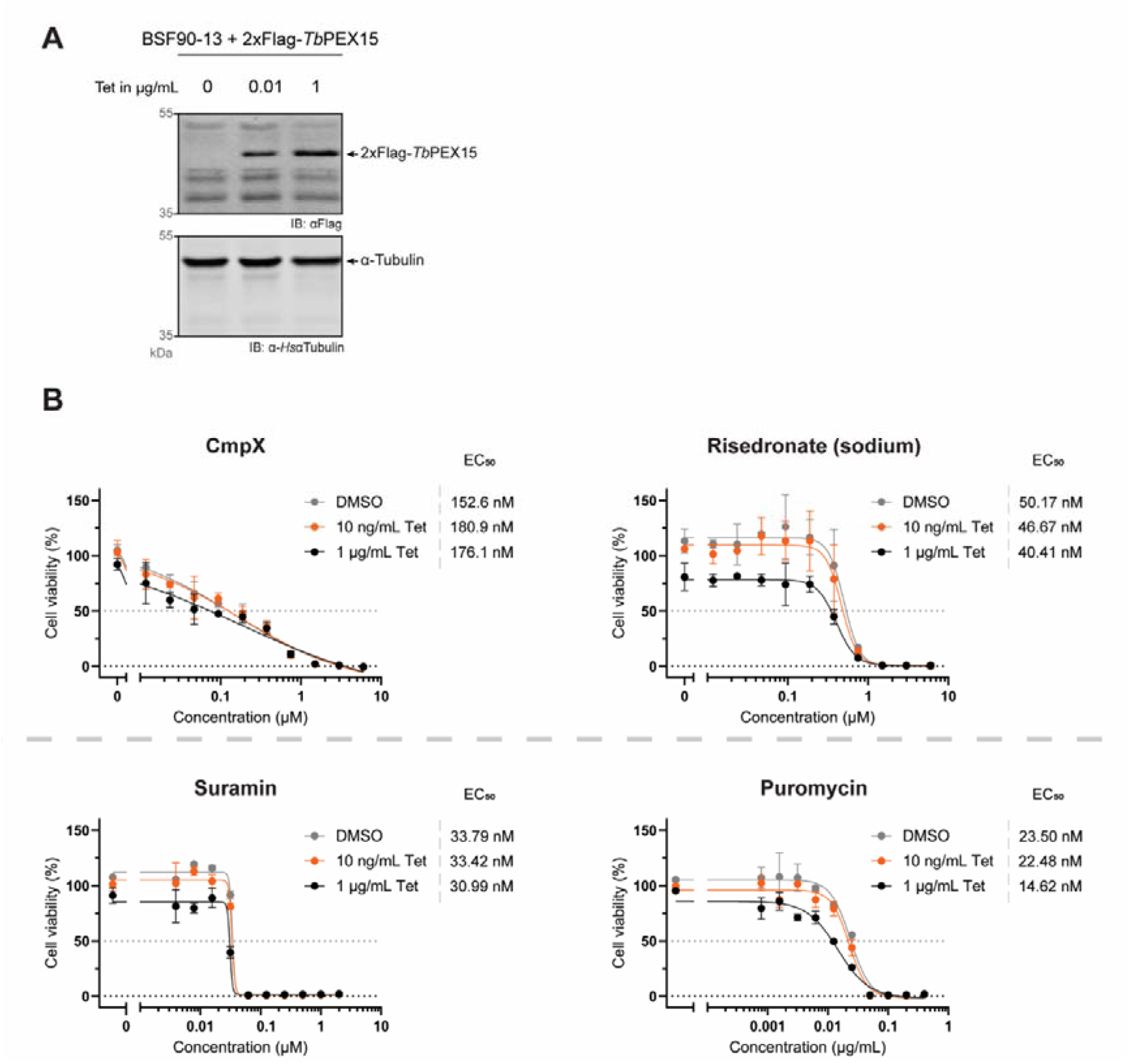
Effect of PEX15 overexpression on drug’s killing effect. (**A**) BSF90-13 were transfected with an overexpression construct coding for 2xFlag-*Tb*PEX15. Induction with 10 ng/mL or 1 μg/mL leads to different rates of protein overexpression. Treatment with DMSO does not lead to expression of 2xFlag-*Tb*PEX15 showing that it can be stringently induced. (**B**) Cells with varying 2xFlag-*Tb*PEX15 overexpression levels were treated with a serial dilution of CmpX (left, top) and risedronate (right, top) or suramin (bottom, left) and puromycin (bottom, right), which were used as controls. Overexpression of 2xFlag-*Tb*PEX15 did not affect the efficacy of the compounds on cell survival. Shown are the results of two biological replicates, error bars indicate ±SD. EC_50_ values were determined via GraphPad Prism 10.0.0.

**Figure S7.**
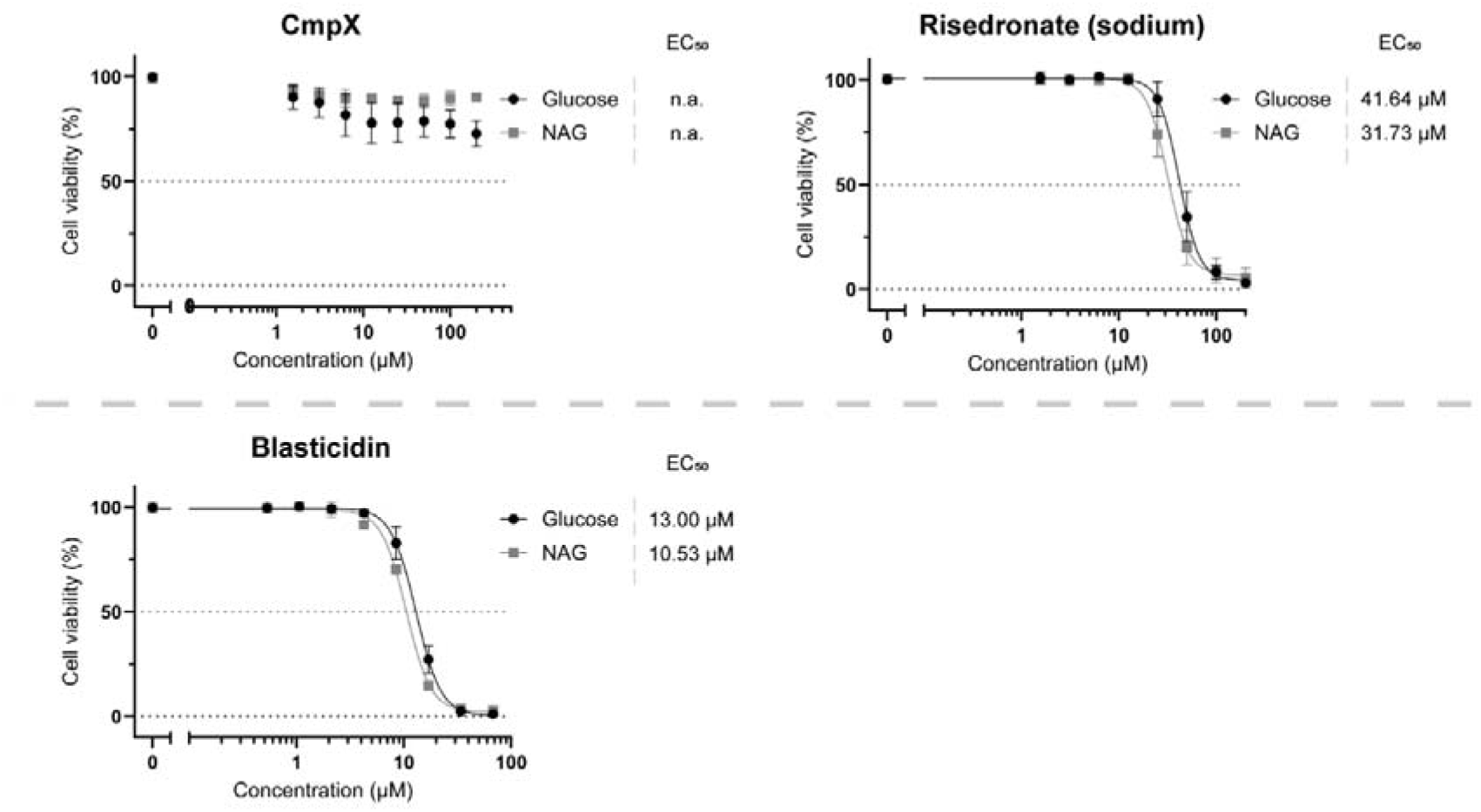
Effect of compounds on procyclic form *T. brucei* cell survival grown in different media. Cells were treated with a two-fold serial dilution of CmpX and risedronate starting from 200 μM or blasticidin as control starting from 68 μM. After 72 h, cell viability was assessed using resazurin. Cells were grown in (1) medium lacking glucose, supplemented with N-acetylglucosamine (NAG, grey data points) to avoid residual uptake of glucose from fetal bovine serum or (2) medium supplemented with glucose (black data points). In presence of glucose, procyclic form (PCF) cells metabolize glucose and thus, depend on functional glycosomes. In the absence of glucose, amino acids are metabolized and disruption of glycosome biogenesis is not lethal. CmpX does not affect survival of PCF cells irrespective of the culture medium. Risedronate kills PCF cells with EC_50_ values of 41.64 μM and 31.73 μM in medium with and without glucose, respectively. Shown are the results of three biological replicates, error bars indicate ±SD. EC_50_ values were determined via GraphPad Prism 10.0.0.

**Figure S8.**
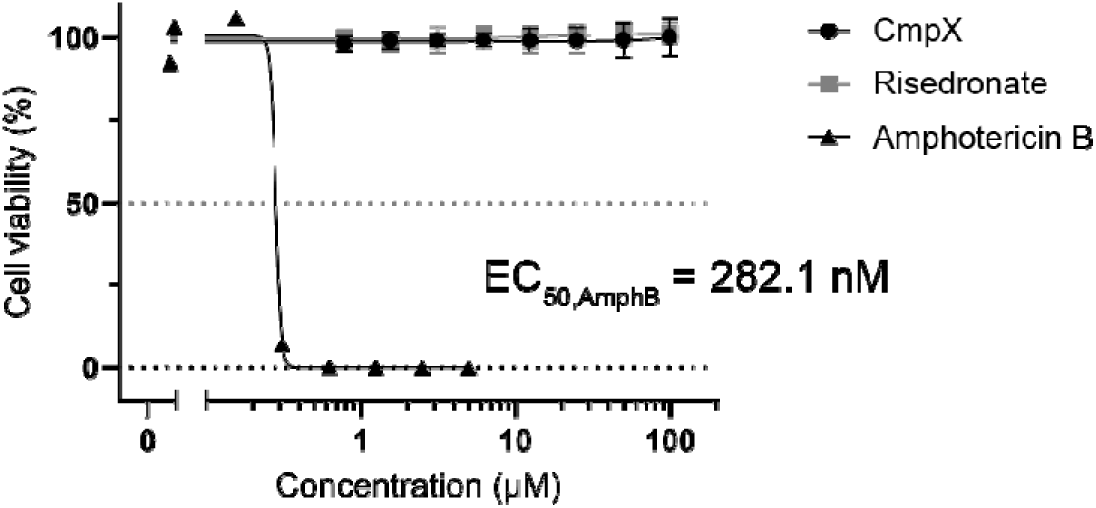
Anti-leishmanial activity of CmpX and risedronate. *Leishmania tarentolae* promastigotes were treated with a two-fold serial dilution of CmpX (black, dots) and risedronate (grey, squares) starting from 100 μM. Amphotericin B (black, triangles) was used as positive control and has an EC_50_ of 282.1 nM. The identified inhibitors are not active against *L. tarentolae* promastigotes.

